# Asymmetric requirement for α-tubulin over β-tubulin

**DOI:** 10.1101/2022.02.17.480930

**Authors:** Linnea C. Wethekam, Jeffrey K. Moore

## Abstract

How cells regulate the supply of α- and β-tubulin monomers to meet the demand for αβ- heterodimers while avoiding consequences of monomer imbalance is not understood. We investigate the role of gene copy number in tubulin regulation and how shifting the expression of α- or β-tubulin genes impacts tubulin proteostasis and microtubule function. We find that α- tubulin gene copy number is important for maintaining an excess α-tubulin protein compared to β-tubulin protein and preventing accumulation of super-stoichiometric β-tubulin. Super-stoichiometric β-tubulin is toxic to cells, leading to loss of microtubules, formation of non-microtubule assemblies of tubulin, and disrupted cell proliferation. In contrast, decreased β- tubulin or increased α-tubulin has minor effects. We provide evidence that cells rapidly equilibrate the concentration of α-tubulin protein during shifts in α-tubulin isotype expression to maintain a ratio in excess of β-tubulin. We propose an asymmetric relationship between α- and β-tubulins, where α-tubulins are maintained in excess to supply αβ-heterodimers and limit the accumulation of β-tubulin monomers.

## Introduction

Cytoskeletal networks are assembled from thousands of protein building blocks; therefore, the size and architecture of these networks sets a demand for the biogenesis and maintenance of these pieces. That demand varies across organisms, cell types within an organism, and even time within a cell. For example, cells of the vertebrate brain require extensive microtubule networks for migration, process formation, and intracellular trafficking. Accordingly, the αβ-tubulin proteins that form microtubules represent approximately 25% of the soluble protein in the mouse brain, compared to 3% of the soluble protein in cultured mouse fibroblasts and less than 1% of soluble protein in the mouse liver (Hiller and Weber, 1978).

Within the lifetime of a cell, the demand for tubulin can shift in response to developmental cues or during cell division. For example, the amount of soluble αβ-tubulin protein in HeLa cells more than doubles from the G1 to mitotic phases of the cell cycle (Bravo and Celis, 1980). These observations raise the fundamental question of how the biogenesis and maintenance of α- and β-tubulin monomers are coordinated to meet the cytoskeletal demands of a cell.

One potential mechanism for meeting different tubulin demands is through the expansion and diversification of α- and β-tubulin gene families, known as ‘isotypes’. Isotypes might serve as transcriptional modules for cells and organisms to meet a tubulin demand, with tissue specific function and expression (Raff, 1984; Kemphues et al., 1982; Latremoliere et al., 2018; Dumontet et al., 1996). Humans have 8-10 α and 7-9 β-tubulin isotypes, and these are expressed at different levels according to cell type and developmental stage (Findeisen et al., 2014; Leandro-García et al., 2010; Park et al., 2021). The expansion of the number of α- and β-tubulin genes provides a variety of transcriptional modules for creating programs of tubulin expression in specific cellular contexts.

However, single-celled organisms also contain multiple copies of tubulin genes. For example, the budding yeast *Saccharomyces cerevisiae* possesses two α-tubulin genes, *TUB1* and *TUB3*, and a single β-tubulin gene, *TUB2.* The two α-tubulins generate approximately equal levels of mRNA, but different levels of soluble protein (Nsamba et al., 2021; Kilmartin and Adams, 1984; Gupta et al., 2002; Gartz Hanson et al., 2016; Barnes et al., 1992). This suggests that families of tubulin genes work additively to supply tubulin, but that the composition of the soluble tubulin pool must also be regulated by additional post-transcriptional mechanisms.

Proteostasis represents a second potential mechanism for meeting tubulin demand.

Here, we consider tubulin proteostasis to consist of the biogenesis of α- and β-tubulin monomers, the equilibrium between monomer and heterodimer states, and the degradation of tubulin. It is well-established that newly synthesized α- or β-tubulin monomers are folded by cytosolic chaperonin and prefoldin, and then assembled into heterodimers by complexing with a series of tubulin binding cofactors (Zabala and Cowan, 1992; Abruzzi *et al*., 2002; Nithianantham *et al*., 2015; TBCs; Figure 1A). The αβ-heterodimer state is not fixed; rather, purified heterodimers can undergo reversible dissociation with moderately fast kinetics into stable monomers of α- and β-tubulin (Montecinos-Franjola et al., 2016). A wide range of dissociation constants have been reported for purified αβ-heterodimers, from 0.1 nM to 1.0 µM (Mejillano and Himes 1989; Detrich and Williams 1978; Caplow and Fee 2002; Fineberg et al., 2020). This apparent disagreement may be attributable to differences in experimental conditions but also to biologically relevant differences in tubulins. αβ-heterodimers purified from different organisms or from different tissues within an organism exhibit dissociation constants that differ by as much as 150-fold in the same experiment (Montecinos-Franjola et al., 2019). Furthermore, there is evidence that cells preferentially sort α- or β-tubulin isotypes into different heterodimer pairs, which could be based on different affinities between α- and β-isotypes (Hoyle et al., 2001). TBCs are good candidates for regulating the monomer-dimer equilibrium, since they can promote subunit exchange in pre-existing heterodimers *in vitro* and are important for maintaining the polymerization-competent pool in cells (Li and Moore, 2020). However, the regulation of monomer-dimer equilibrium in cells is largely uncharacterized. Tubulin turnover is also poorly understood. We know tubulin is degraded by the proteasome (Huff et al., 2010), but whether the heterodimer or monomer state is preferentially targeted for degradation is an open question. In general, tubulin proteostasis is likely to play an important role in meeting tubulin demand but is unexplored in the field.

**Figure 1:**
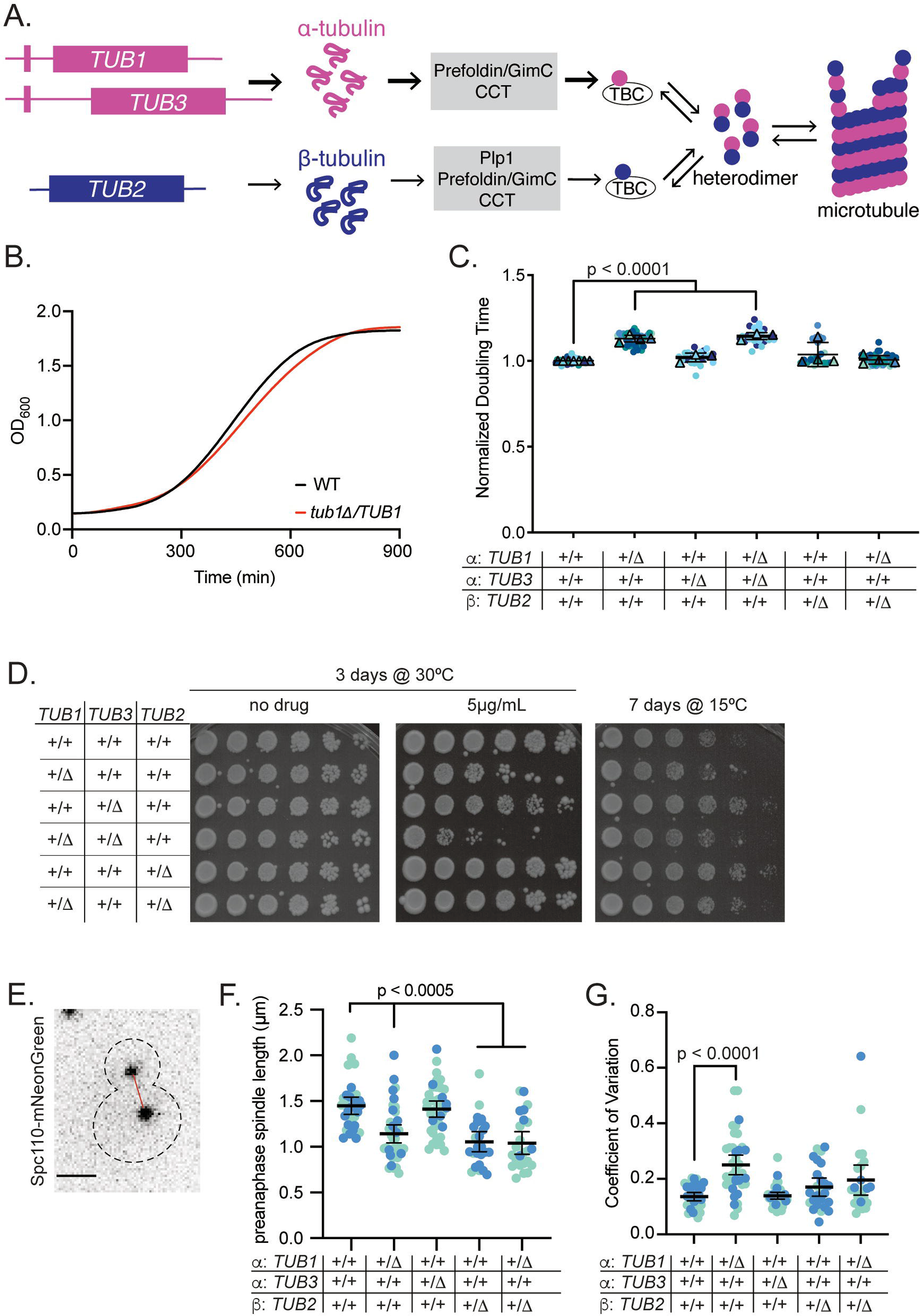
Distinct requirements for α- and β-tubulin gene copy number in microtubule function A. Current model for how cells build microtubules from tubulin genes through folding and assembly of polymerization competent heterodimers. B. Representative growth curves for wild-type and *TUB1/tub1*Δ heterozygous cells. All cultures were grown at 30°C with agitation for 24 hours, and OD600 was measured every 5 minutes. C. Doubling times of indicated heterozygous tubulin mutants normalized to wild-type controls. For each genotype, at least four technical replicates of two biological replicates were utilized across at least three independent experiments. Circles represent cultures in separate wells and triangles are the means of each day. Colors indicate independent experiments. P-value between wild type and mutant based on t-test after one-way ANOVA with a Tukey post-hoc test. Bars are mean ± 95% CI. D. Ten-fold dilution series of indicated strains were spotted onto rich media or rich media supplemented with benomyl. Cells were grown at the indicated temperature for the indicated number of days. E. Representative image of a wild-type pre-anaphase cell expressing Spc110-mNeonGreen. Red line indicates distance between two poles. Scalebar = 1 µm. Image is a maximum intensity projection of z series. F. Mean pre-anaphase spindle length per cell across 5-minute imaging timeframe. For each genotype, two biological replicates were used across two independent experiments. Each dot represents the mean from one cell. Colors indicate independent experiments. P-value between wild type and mutant based on t-test after one-way ANOVA. Bars are mean ± 95% CI. G. Coefficient of variance of pre-anaphase spindle length per cell across 5-minute imaging timeframe, from the dataset used in 1F. Each dot represents the mean from one cell. Colors indicate independent experiments. P-value between wild type and mutant based on t-test after one-way ANOVA. Bars are mean ± 95% CI.

In this study we sought to better understand how cells coordinate α- and β-tubulin across genes and protein to meet the demand for functional αβ-heterodimers. We used the budding yeast model due to its simplified repertoire of α- and β-tubulins, genetic tractability, and well-defined microtubule networks. We find that cells maintain an excess of α-tubulin compared to β- tubulin and are more sensitive to loss of α-tubulin genes than β-tubulin genes in diploid cells.

Removing a copy of the α-tubulin isotype *TUB1* causes slower proliferation, increased sensitivity to microtubule stress, and unstable mitotic spindles. We also find that super-stoichiometric levels of α- or β-tubulin create non-microtubule assemblies of tubulin, but only super-stoichiometric β-tubulin disrupts the microtubule cytoskeleton. In contrast, when α-tubulin is overexpressed, levels of other α-tubulin isotypes decrease in response and ultimately return cells to endogenous levels of α-tubulin. We propose a model where cells use isotypes to create an excess of α-tubulin expression, and then rapidly exchange α-tubulin protein to ensure sufficient heterodimer production and prevent the accumulation of β-tubulin monomers.

## Results

### Distinct requirements for α- and β-tubulin gene copy number in microtubule function

We used three experiments to test the prediction that altering gene copy number disrupts microtubule function. We first compared the proliferation of a panel of diploid yeast strains in which we knocked out one copy of an α- or β-tubulin gene. We grew the cells in liquid media, calculated doubling times during log phase growth, and normalized our data to wild-type control cells, which take approximately 127 ± 3 minutes to double in our assay (Figure 1B). We find that diploid cells with only one copy of the α-tubulin isotype *TUB1* exhibit doubling times that are 13.1% longer than wild type (140 ± 6 minutes, p <0.0001; Figure 1B,C). In contrast, diploid cells with only one copy of the α-tubulin isotype *TUB3* have doubling times that are indistinguishable from wild type (Figure 1C, p = 0.9470). Diploid cells that only have one copy each of *TUB1* and *TUB3* exhibit doubling times that are 14.5% longer than wild type, similar to *TUB1* single heterozygous nulls (149 ± 4 minutes, p <0.0001; Figure 1C). Removing one copy of β-tubulin did not elicit a growth phenotype; cells with only one copy of *TUB2* exhibit a doubling time similar to wild type. Interestingly, double mutants lacking one copy each of *TUB1* and *TUB2* were also similar to wild-type controls (Figure 1C, p = 0.4733). This suggests that cell fitness is more sensitive to the gene copy number of α-tubulin, specifically *TUB1*, than β-tubulin, and that sensitivity to α-tubulin depletion is rescued by simultaneous depletion of β-tubulin.

For our second test of tubulin function, we compared sensitivity to two different microtubule stressors – the destabilizing drug, benomyl, and low temperatures. We plated serial dilutions of the panel of tubulin heterozygote strains on rich media or rich media containing benomyl, and incubated the plates either at the standard temperature for culturing budding yeast (30°C) or at a lower temperature where yeast grow slowly and exhibit hypersensitivity to disruption of microtubule function (15°C; Li and Moore, 2020). We find that the heterozygous mutants that exhibit slower doubling times above are also more sensitive to benomyl and low temperatures (Figure 1D). These results also indicate a stronger requirement for α-tubulin gene copy number than β-tubulin gene copy number. Since the *TUB1 TUB3* double heterozygous mutants exhibit a phenotype similar to that of the *TUB1* single heterozygote, we focused our subsequent experiments on the *TUB1* or *TUB3* single heterozygotes.

For our third test, we measured the lengths of pre-anaphase spindles, which are formed by interdigitating microtubules emanating from the two spindle pole bodies (SPBs) (Winey et al., 1995). We tracked Spc110-mNeonGreen marked SPBs in asynchronous, living cells over five minutes and identified pre-anaphase spindles as those that did not exhibit sustained lengths beyond 2.2 µm during our imaging period. We then measured pole-to-pole distance in 3 dimensions (X,Y,Z) to determine the average spindle length and length variation over time within the spindle of each cell (Thomas et al., 2020; Figure 1E-G). Diploid cells with only one copy of *TUB1* exhibit shorter pre-anaphase spindles compared to wild-type controls (Figure 1F, p <0.0001). In contrast, diploid cells with one copy of *TUB3* have mean preanaphase spindles that are a similar length to wild type (Figure 1F, p = 0.9838). Diploid cells with one copy of *TUB2* also exhibit shorter pre-anaphase spindles than wild-type controls (Figure 1F, p < 0.0001). Cells lacking one copy of *TUB1* and *TUB2* together exhibit shorter pre-anaphase spindles that are not significantly different from either single heterozygote (Figure 1F, p < 0.0001 to WT, p > 0.63 to *TUB1* or *TUB2* alone). Since we tracked spindle length over time we were able to measure the stability of each spindle by calculating the coefficient of variation. Whereas pre-anaphase spindles in wild-type cells exhibit only small variation in spindle length over time, cells lacking one copy of *TUB1* exhibit increased coefficients, indicating that spindle length is more variable over time in these cells compared to wild type (Figure 1G, p < 0.0001). In contrast, diploid cells lacking a copy of either *TUB3* or *TUB2* have coefficient of variation similar to wild type (Figure 1G, p > 0.5446). Furthermore, cells lacking one copy of both *TUB1* and *TUB2* together also show a coefficient of variation not different from wild type (Figure 1G, p = 0.0684). To summarize these results, loss of either α- or β-tubulin genes results in shorter spindles, but loss of *TUB1* uniquely results in unstable spindles. We conclude that budding yeast is more sensitive to loss of α-tubulin genes than β-tubulin genes, and that this effect may be attributable to the creation of excess β-tubulin expression.

### α- and β-tubulin gene copy number determines polymerization activity and the balance between subunits

We next investigated how altering tubulin gene copy number affects microtubule polymerization and tubulin protein levels. We predicted that decreasing gene copy number for either α- or β- tubulin would decrease heterodimer availability in cells, leading to shorter microtubules and slower polymerization rates. To test this prediction, we measured the dynamics of individual astral microtubules over time using CLIP-170/Bik1-3GFP to label growing microtubule ends (Figure 2A, B). Since there was no difference in pre-anaphase spindle length between WT and the *TUB3* heterozygous null we did not include this strain in this experiment. By comparing the full dataset of length measurements at each timepoint we find that *TUB1* and *TUB2* single heterozygous mutants and *TUB1 TUB2* double heterozygous mutants each exhibit shorter astral microtubules that sample a narrower range of lengths than the wild-type controls (Figure 2C). The length distributions for the mutants are all similar to each other, with the exception that *TUB1* heterozygotes have some microtubules that reach long lengths (Figure 2C). In addition, polymerization rate is decreased in each of the mutant strains in our panel (Figure 2D; see full set of microtubule dynamics measurements in Table 1). Wild-type diploids have a median polymerization rate of 1.271 µm/min whereas removing one copy of *TUB1* alone decreased polymerization rate to 1.146 µm/min (Table 1, Figure 2D). Loss of *TUB2* alone or the combined loss of *TUB1* and *TUB2* showed even slower polymerization rates (median = 1.023 µm/min and 0.9697 µm/min respectively, Figure 2D). We conclude that gene copy number for both α- and β- tubulin is important for heterodimer activity, but the lower rates observed in *TUB2* heterozygotes suggest that β-tubulin may be limiting for polymerization rate.

**Figure 2:**
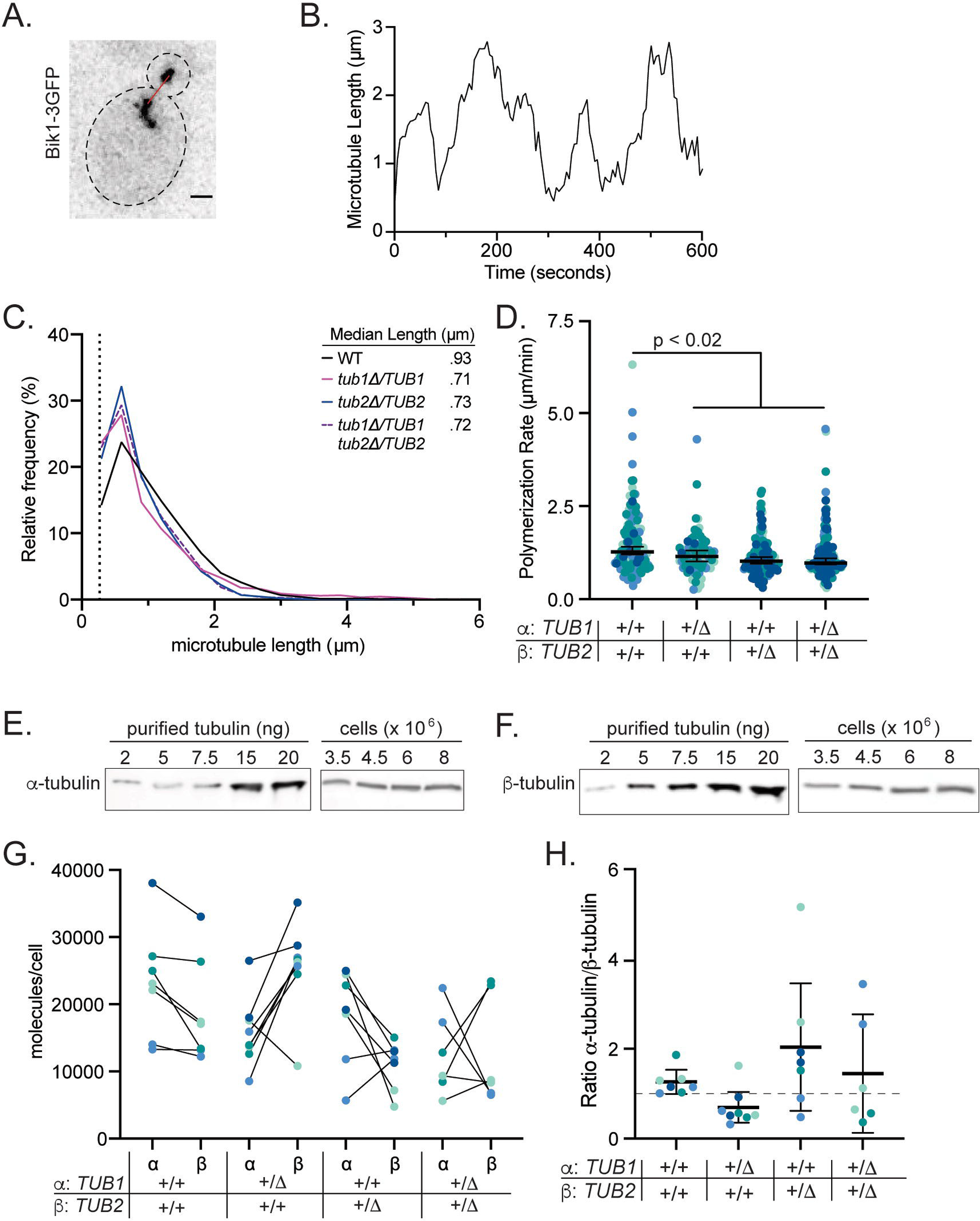
Changes in gene copy number alter microtubule activity A. Representative image of a wild-type cell expressing Bik1-3GFP. Red line indicates measured length between plus-end and spindle pole body. Scalebar = 1 µm. mage is a maximum intensity projection of z series. B. Life plot of a single microtubule from a wild-type cell. Microtubule lengths were measured at 5-second intervals. C. Histogram of all astral microtubule lengths from time lapse imaging of wild type, *TUB1/tub1Δ*, *TUB2/tub2Δ*, and *TUB1/tub1Δ TUB2/tub2Δ*. Data are from at least three separate experiments for each genotype, and a total of at least 20 cells were analyzed for each genotype. D. Polymerization rates of astral microtubules. Each dot represents a single polymerization event and dots are colored by experimental day. P-value between wild type and mutant based on t-test after one-way ANOVA. Bars are median ± 95% CI. E. Representative image of quantitative western blot of α-tubulin in wild-type cells to determine molecules per cell. Known mass of purified yeast tubulin was used to make a standard curve. F. Representative image of quantitative western blot of β-tubulin in wild-type cells to determine molecules per cell. Known mass of purified yeast tubulin was used to make a standard curve. G. Paired molecules of α- or β-tubulin per cell for each genotype. Protein mass (ng) was converted to molecules per cell. Data represent three to four independent experiments with two biological replicates and each dot is mean molecules per cell of the four dilutions. Dots are colored by experiment. H. Ratio of α- to β-tubulin in cells with indicated genotypes. Ratios were calculated from data in 2G. Bars are mean ± 95% CI.

**TABLE 1:** Astral microtubule dynamics and soluble tubulin levels in diploid cells. Values in bold have p <0.05 from wild type, based on a T-test of p <0.05 from an ANOVA with a Tukey post- hoc test. a. Median values (95% CI) b. Mean values ± 95% CI c. Mean values ± 95% CI; values from Figure 2G

| Genotype | Median length<br>($\mu\text{m}$)<sup>a</sup> | Polym Rate<br>($\mu\text{m}/\text{min}$)<sup>a</sup> | Depolym Rate<br>($\mu\text{m}/\text{min}$)<sup>a</sup> | Catastrophe<br>frequency<br>(events/min)<sup>b</sup> | Rescue<br>frequency<br>(events/min)<sup>b</sup> | Dynamicity<br>(subunits/s)<sup>b</sup> | α-tubulin<br>molecules<br>per cell<sup>c</sup> | β-tubulin<br>molecules<br>per cell<sup>c</sup> |
| --- | --- | --- | --- | --- | --- | --- | --- | --- |
| Wild type<br>(n = 30 cells) | 0.93<br>(0.91 – 0.96) | 1.27<br>(1.2 – 1.41) | 1.91<br>(1.72 – 2.22) | 0.81 ± 0.11 | 1.02 ± 0.19 | 0.49 ± 0.03 | 2.3x10<sup>4</sup> ± 0.7 | 1.9x10<sup>4</sup> ± 0.7 |
| *tub1Δ/TUB1*<br>(n = 20 cells) | **0.71**<br>**(0.69 – 0.73)** | **1.15**<br>**(1.02 – 1.31)** | **1.60**<br>**(1.34 – 1.84)** | 0.62 ± 0.12 | 0.9 ± 0.2 | 0.43 ± 0.06 | **1.6x10<sup>4</sup> ± 0.4** | 2.6x10<sup>4</sup> ± 0.6 |
| *tub2Δ/TUB2*<br>(n = 35 cells) | **0.73**<br>**(0.7 – 0.76)** | **1.02**<br>**(0.95 – 1.14)** | **1.53**<br>**(1.42 – 1.71)** | 0.8 ± 0.09 | 0.87 ± 0.12 | **0.41 ± 0.02** | 1.8x10<sup>4</sup> ± 0.7 | **1.1x10<sup>4</sup> ± 0.3** |
| *tub1Δ/TUB1*<br>*tub2Δ/TUB2*<br>(n = 36 cells) | **0.72**<br>**(0.7 – 0.74)** | **0.97**<br>**(0.93 – 1.1)** | **1.43**<br>**(1.27 – 1.70)** | 0.74 ± 0.11 | 0.88 ± 0.14 | **0.39 ± 0.04** | **1.3x10<sup>4</sup> ± 0.7** | 1.3x10<sup>4</sup> ± 0.8 |

Next, we measured the levels of soluble α- and β-tubulin in cells with altered gene copy number. We first created standard curves of purified yeast tubulin on a western blot probed with antibodies for α- and β-tubulin (Figure 2E, F; S1; see Materials and Methods). We then compared western blots of protein lysates from log-phase cells, creating dilution series that we matched to the number of cells in the culture, and used the standard curves to calculate the molecules of soluble α- or β-tubulin per cell. We find that levels of soluble α-tubulin are higher than levels of soluble β-tubulin in wild-type control cells, with a mean ratio of 1.26 α-tubulins per β-tubulin (Figure 2G, H). This excess of α-tubulin was observed across genotypes, with one exception – *TUB1* heterozygotes exhibit higher levels of β-tubulin than α-tubulin, with a mean ratio of 0.70 α-tubulins per β-tubulin (Figure 2G and H). These results indicate that wild-type cells normally contain an excess of α-tubulin and that lowering α-tubulin gene copy number in *TUB1* heterozygotes creates an aberrant excess of β-tubulin.

### Super-stoichiometric β-tubulin creates aberrant tubulin assemblies

Our results thus far suggest that the unique phenotypes of *TUB1* heterozygous cells may be due to the creation of super-stoichiometric β-tubulin. We next sought to directly test how increasing β-tubulin expression impacts fitness and microtubule function. Previous studies have found that constitutive expression of an additional copy of *TUB2* is lethal in budding yeast (Weinstein and Solomon, 1990; Burke et al.,1989). We therefore designed a plasmid-based tool for conditionally overexpressing ectopic *TUB2* from a galactose-inducible promoter in wild-type cells. By collecting cells at timepoints after adding galactose and performing western blotting, we determined relative α- or β-tubulin levels during the time course of ectopic β-tubulin expression (Figure 3A, B). After 15 minutes of galactose induction, the level of β-tubulin in soluble protein lysates is 1.3x the amount measured in uninduced control cells (Figure 3B, C). Cells collected after 15 minutes of induction and then plated to glucose-containing media to shut off ectopic β-tubulin expression show a strong inhibition of colony formation (Figure 3D; Weinstein and Solomon, 1990). At one hour of galactose induction, β-tubulin is increased to 2x the level measured in uninduced control cells and nearly all cells fail to form colonies (Figure 3B-D). At no point during the time course of β-tubulin overexpression did we observe any increase in α-tubulin levels (Figure 3B, C). This result demonstrates that even a small excess of β-tubulin by itself is acutely toxic to cells.

**Figure 3:**
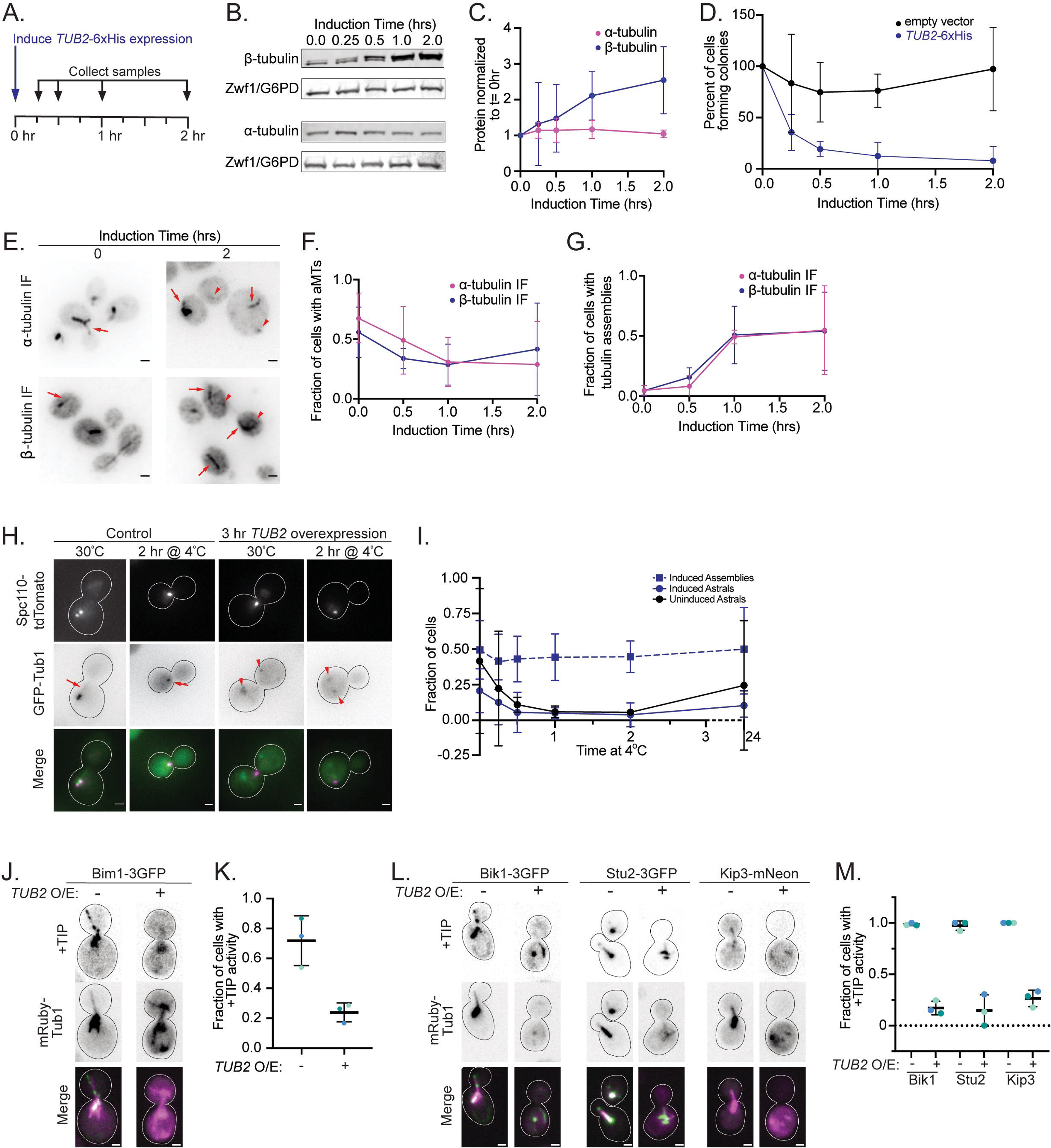
Super-stoichiometric β-tubulin creates aberrant tubulin assemblies A. Schematic for β-tubulin induction experiment. B. Representative western blot of β- and α-tubulin during *TUB2-6xHIS* induction. Blots were probed for α- or β-tubulin, and Zwf1 (G6PD) as a loading control. C. Quantification of *TUB2-6xHis* induction for both α- and β-tubulin across overexpression. Dots represent mean ± SD from four independent induction experiments. D. Quantification of cells forming colonies after *TUB2-6xHis* induction or empty vector control. Dots represent mean ± SD from four independent induction experiments. E. Example images of wild-type cells overexpressing β-tubulin for 0 or 2 hours and stained for α- or β-tubulin. Arrows indicate astral microtubules; arrowheads indicate tubulin assemblies. Scale bar = 1 µm. Images are maximum intensity projections of in focus z positions. F. Quantification of astral microtubules (aMTs) across induction time for cells stained for either α- or β-tubulin. Dots represent mean ± 95% CI from three independent experiments with at least 100 cells in each experiment. G. Quantification of tubulin assemblies across induction time for cells stained for either α- and β-tubulin. Dots represent mean ± 95% CI from three independent experiments with at least 100 cells in each experiment. H. Example images of cells expressing Spc110-tdTomato and GFP-Tub1. *TUB2-6xHis* was induced for 3 hours and then cells were shifted to 4°C for 2 hours to depolymerize microtubules. Uninduced control cells are also shown. Arrows indicate microtubules; arrowheads indicate tubulin assemblies. Scale bar = 1 µm. Images are maximum intensity projections of z series. I. Quantification of the fraction of astral microtubules or tubulin assemblies in induced or uninduced cells after indicated time at 4°C. Dots are mean ± 95% CI from at least 100 cells in three independent experiments. J. Example images of cells expressing Bim1-3GFP and mRuby-Tub1 with *TUB2-6xHis* induced or uninduced for 2 hours. Confocal images were processed in Fiji using the Despeckle filter and stacks were converted to sum projections. Scale bar = 1 µm. K. Quantification of the fraction of cells exhibiting a focus of Bim1-GFP that tracks a dynamic microtubule plus end in uninduced or induced cells. Each dot represents a single experiment. Bars are mean ± 95% CI. L. Example images of cells expressing the indicated +TIP and mRuby-Tub1 with *TUB2- 6xHis* induced or uninduced for 2 hours. Confocal images were processed in Fiji using the Despeckle filter and stacks were converted to sum projections. Scale bar = 1 µm. M. Quantification of the fraction of cells exhibiting a focus of +TIP that tracks a dynamic microtubule plus end in uninduced or induced cells. Each dot represents a single experiment. Bars are mean ± 95% CI.

This result led us to test how β-tubulin overexpression alters the microtubule cytoskeleton. We first used immunofluorescence to visualize α- and β-tubulin. We expect microtubules containing a 1:1 stoichiometry of α- and β-tubulin would exhibit a characteristic morphology and abundance when stained with antibodies to either tubulin. In wild-type haploid cells, we found that the level of β-tubulin overexpression (0.5, 1, or 2 hours, which represent 1.5x, 2x and 2.5x the level of β-tubulin in uninduced control cells; Figure 3C) was inversely proportional to the presence of astral and nuclear microtubules (Figure 3E). Staining for α- tubulin shows that the frequency of cells containing astral microtubules steadily decreases as the level of β-tubulin overexpression increases (Figure 3F). Staining for β-tubulin shows a similar trend, although the images were less clear due to increasing background signal at greater levels of β-tubulin expression and generally poorer staining from the β-tubulin antibody (Figure 3E-F). While microtubules are lost during β-tubulin overexpression, we observed the formation of alternative structures that stained with tubulin antibodies. These structures could be distinguished from microtubules because they are typically disconnected from the SPBs and/or orthogonal to microtubules in the same cell, and are heterogeneous in shape and size, from small foci to tangled filaments (Figure 3E, G; S2). We collectively termed these structures ‘tubulin assemblies’. We scored cells for the appearance of these assemblies and found that they first emerged after one hour of β-tubulin overexpression, and by two hours 50% of cells had at least one tubulin assembly (Figure 3E, G). Similar tubulin assemblies could be detected by β- tubulin or α-tubulin immunofluorescence, indicating that tubulin assemblies contain both tubulins. These results suggest that super-stoichiometric β-tubulin dominantly disrupts normal microtubule architecture and creates new α- and β-tubulin-containing assemblies.

To determine whether tubulin assemblies formed in the presence of super-stoichiometric β-tubulin exhibit properties that are distinct from microtubules, we first asked whether they are cold labile. We tested this by overexpressing β-tubulin in cells with labeled spindle pole bodies (Spc110-tdTomato) and a GFP fusion to the N-terminus of ectopically expressed α-tubulin (GFP-Tub1). Cells were induced to overexpress β-tubulin for 3 hours, then shifted to 4°C for 0.25, 0.5, 1, 2, and 24 hours, fixed, and imaged (Figure 3H). Whereas microtubules are lost in uninduced control cells within 1 hour of the shift to 4°C, tubulin assemblies are retained in cells overexpressing β-tubulin for many hours after the shift to 4°C (Figure 3I). These assemblies appeared as a combination of linear filaments and non-linear clusters that contained GFP-Tub1 and were not connected to the spindle pole bodies (Figure 3H). The characteristics of cold resistance and dissociation from the spindle pole bodies suggest that β-tubulin overexpression creates tubulin assemblies that are not bona fide microtubules.

As a second test, we asked whether the tubulin assemblies formed during β-tubulin overexpression recruit microtubule-associated proteins (MAPs). We tested this using four well-characterized MAPs that are known to use distinct modes of binding to microtubule plus ends. We first tested the plus-end binding protein Bim1, the budding yeast member of the EB protein family (Tirnauer and Bierer, 2000). EB proteins bind specifically to the microtubule lattice via a binding site that consists of α- and β- tubulins from four adjacent heterodimers (Maurer et al., 2012). Bim1 selectively binds to a transition state of GTP hydrolysis that accompanies microtubule polymerization (Howes et al., 2018). Therefore, Bim1 localization provides a marker for a specific microtubule lattice state and would not be expected to bind to a tubulin assembly that lacks these properties. Cells expressing Bim1-3GFP from its native locus and mRuby-Tub1 to label tubulin were induced to overexpress β-tubulin for two hours and then living cells were imaged by time lapse confocal microscopy (Figure 3J). 75% of uninduced control cells exhibit at least one focus of Bim1-3GFP in the cytoplasm (Figure 3K). These foci localize to the end of microtubules labeled with mRuby-Tub1, and track plus ends as the microtubules polymerize and depolymerize over time. In most cells overexpressing β-tubulin, Bim1-3GFP signal is diffuse in the cytoplasm with no accumulation on tubulin assemblies (Figure 3J), while 23% of cells show Bim1-3GFP localization to an mRuby-Tub1 labeled astral microtubule (Figure 3K). This suggests that tubulin assemblies formed during β-tubulin overexpression do not contain the lattice state that is normally found at the microtubule plus end.

To further interrogate the composition of the tubulin assemblies we completed similar experiments using three other plus-end binding proteins: Bik1/CLIP-170, Stu2/XMAP215, and Kip3/kinesin-8. Bik1-3GFP localizes to microtubules via CAP-Gly domains that bind to EEY/F motifs in α-tubulins (Pierre et al., 1992; Weisbrich et al., 2007; Badin-Larçon et al., 2004). We predicted that if the tubulin assemblies contain sub-stoichiometric levels of α-tubulin due to excess β-tubulin, compared to the 1:1 stoichiometry of α- and β in microtubules, then Bik1 will exhibit diminished localization to tubulin assemblies. Indeed, cells overexpressing β-tubulin exhibit either diffuse Bik1-3GFP signal in the cytoplasm and/or colocalization along tubulin assemblies without clear enrichment at filament ends (Figure 3L, M). This suggests that β- tubulin induced assemblies do contain α-tubulin but lack the plus end that is normally recognized by Bik1.

Stu2/XMAP215 uses a combination of αβ-heterodimer binding by its TOG domains and a poorly defined lattice-binding activity by its basic domain to localize to microtubule plus ends (Ayaz et al., 2012; Geyer et al., 2018). While the plus-end localization of Stu2-3GFP is lost when β-tubulin is overexpressed, we find that Stu2-3GFP exhibits strong co-localization to tubulin assemblies, albeit without enrichment at filament ends (Figure 3L, M). This indicates that tubulin assemblies may contain strong binding sites for either the TOG domains or the basic domain of Stu2.

Finally, we examined the plus-end directed kinesin-8, Kip3, which binds to microtubules at the intradimer interface and walks towards the plus end where it induces microtubule depolymerization (Varga *et al*., 2009; Arellano-Santoyo *et al*., 2021 *Preprint*). Kip3-mNeonGreen does not localize to tubulin assemblies, and instead is diffusely localized in the cytoplasm when β-tubulin is overexpressed (Figure 3L, M). This suggests that tubulin assemblies do not contain the intradimer interface in high abundance. Taken together, our results suggest that super-stoichiometric β-tubulin leads to loss of microtubules and the formation of cold-stable tubulin assemblies that lack conventional binding sites for MAPs.

### Cells tolerate super-stoichiometric α-tubulin

If β-tubulin overexpression disrupts the microtubule cytoskeleton, we asked whether overexpressing α-tubulin elicits similar effects. In contrast to β-tubulin, ectopic copies of α- tubulin genes under control of the endogenous promoter are tolerated by budding yeast (Katz, Weinstein, and Solomon 1990). We find that an additional copy of *TUB1* on a low-copy, centromere-containing plasmid increases doubling time by approximately 5% over that observed in wild-type diploid cells (Figure S3A,B). The same *TUB1* plasmid partially rescues the growth defect of heterozygous cells lacking one chromosomal copy of *TUB1*, indicating that the plasmid-borne copy does provide functional *TUB1* (Figure S3B). In other assays of tubulin function, an additional copy of *TUB1* confers strong benomyl resistance and slightly increases the frequency of long microtubules in wild-type diploid cells but did not noticeably alter polymerization rate (Figure S3C-E). When we perform quantitative western blotting, we find that the additional copy of *TUB1* does not change the soluble pool of α- or β-tubulin (Figure S3F; Schatz et al., 1986). Together these data suggest that increasing α-tubulin gene copy number confers resistance to microtubule destabilizing drugs, but otherwise does not lead to major changes in microtubule function or soluble tubulin levels.

We next investigated how cells respond to acute overexpression of *TUB1*, using a galactose-inducible system similar to what we used for *TUB2.* After 15 minutes of galactose induction, the level of α-tubulin in soluble protein lysates is 1.3x the amount measured in uninduced control cells (Figure 4A, B), which is similar to the rate of β-tubulin induction measured in Figure 3. However, we saw no change in the ability of cells to form colonies when α-tubulin is overexpressed (Figure 4C). At one hour of galactose induction, the level of α-tubulin in soluble protein lysates is 1.7x the amount measured in uninduced control cells, and we find no loss in the ability of cells to form colonies (Figure 4B, C). This suggests that excess α-tubulin does not impair fitness, which is in contrast to excess β-tubulin (Figure 3B). To test whether the toxicity associated with β-tubulin overexpression is attributable to increased tubulin levels or to super-stoichiometric β-tubulin, we increased α- and β-tubulin levels while maintaining stoichiometry by simultaneously inducing α- and β-tubulin overexpression from separate plasmids in the same cells (Figure S4A, B). Under these conditions, we find that α- and β-tubulin levels increase with kinetics similar to the individual overexpression experiments, but there is no change in colony formation over our time course of induction (Figure S4C). We conclude that super-stoichiometric β-tubulin is uniquely toxic to cells.

**Figure 4:**
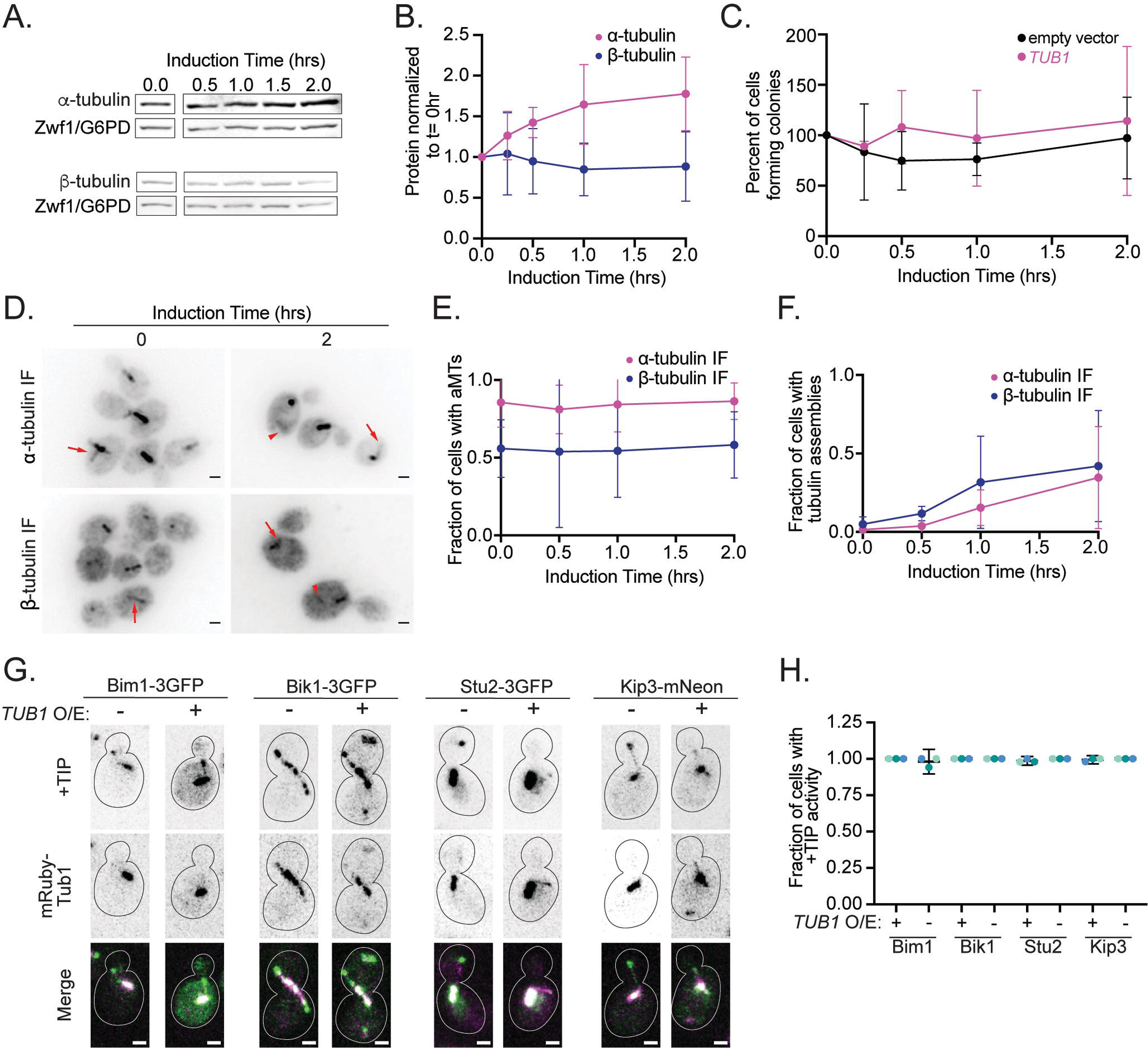
Cells tolerate super-stoichiometric α-tubulin A. Representative western blot of α- and β-tubulin during *TUB1* induction. Blots were probed for α- or β-tubulin and Zwf1 (G6PD) as a loading control. B. Quantification of *TUB1* induction for both α- and β-tubulin across overexpression. Dots represent mean ± SD from four independent induction experiments. C. Quantification of cells forming colonies after *TUB1* or empty vector induction. Dots represent mean ± SD from three independent induction experiments. D. Example images of wild-type cells overexpressing *TUB1* for 0 or 2 hours and stained for α- or β-tubulin. Arrows indicate astral microtubules; arrowheads indicate tubulin assemblies. Scale bar = 1 µm. Images are maximum intensity projections of in focus z positions. E. Quantification of astral microtubules across induction time for both α- and β-tubulin staining. Dots represent mean ± 95% CI from three independent experiments with at least 100 cells per experiment. F. Quantification of tubulin assemblies across induction time for both α- and β-tubulin staining. Dots represent mean ± 95% CI from three independent experiments with at least 100 cells per experiment. G. Example images of cells expressing the indicated +TIP and mRuby-Tub1 with *TUB1* induced or uninduced for 2 hours. Confocal images were processed in Fiji using the Despeckle filter and stacks were converted to sum projections. Scale bar = 1 µm. H. Quantification of the fraction of cells exhibiting a focus of +TIP that tracks a dynamic microtubule plus end in uninduced or induced cells. Each dot represents a single experiment. Bars are mean ± 95% CI.

We next tested whether α-tubulin overexpression alters the microtubule cytoskeleton. We used immunofluorescence to visualize α- and β-tubulin in wild-type haploid cells at the same induction timepoints as our above experiments with β-tubulin. We found no change in the fraction of cells with astral microtubules during α-tubulin overexpression (Figure 4D, E; S5). We also saw tubulin assemblies form during α-tubulin induction, though these always appeared as foci compared to the assemblies formed during β-tubulin overexpression (Figure 4F; S5). This suggests that super-stoichiometric levels of α-tubulin do not disrupt the microtubule cytoskeleton, in contrast to what we find for super-stoichiometric levels of β-tubulin.

We utilized the same panel of plus-end binding proteins to investigate the impact of α- tubulin overexpression on microtubules in living cells. We find that α-tubulin overexpression does not noticeably disrupt microtubule architecture or the localization of the +TIPs Bim1, Bik1, Stu2 and Kip3, even in cells that also exhibit tubulin assemblies labeled with mRuby-Tub1 (Figure 4G,H). Bim1 and Stu2 do not localize to the assemblies formed during α-tubulin overexpression; however, Bik1 and Kip3 do show some localization to these assemblies, albeit at weaker signal intensities than what is observed at microtubule plus ends (Figure 4G, H).

Overall, our results suggest that overexpressed α-tubulin does not interfere with the microtubule cytoskeleton and can form ectopic tubulin assemblies, although the morphology is different than what we observe during β-tubulin overexpression. Together these data indicate that overexpressed α-tubulin is not toxic to cells.

### α-tubulin isotypes Tub1 and Tub3 are balanced to prevent super-stoichiometric α-tubulin levels

Our results thus far suggest a major difference between α- and β-tubulin proteostasis: when α- tubulin expression is increased cells can readily equilibrate the newly expressed protein with existing tubulin to maintain the pool of αβ-tubulin heterodimers in the cell, but increasing β- tubulin expression destroys microtubules and creates a toxic accumulation of β-tubulin protein (Figure 2G; 3). This led us to investigate how the two α-tubulin isotypes in budding yeast, *TUB1* and *TUB3*, might be coordinated to maintain α-tubulin levels. To do this we built a strain in which we fused GFP to the 5’ end of chromosomal *TUB3* and replaced the endogenous *TUB3* promoter with a galactose-inducible promoter from the *GAL1* gene (Figure 5A). We induced GFP-Tub3 expression by adding galactose to log-phase cultures and took samples for microscopy or western blotting at 2, 3 and 24 hours post induction (Figure 5B). The GFP fusion allows us to visualize the GFP-Tub3 production and the rate of assembly into microtubules in cells, and allows us to separate GFP-Tub3 from endogenous untagged Tub1 on a western blot probed for α-tubulin (Figure 5B, E). We first used microscopy to establish the temporal order of GFP signal accumulation in microtubules vs the cytoplasm vs non-microtubule tubulin assemblies (Figure 5C, D). We did this experiment in two ways: first, we induced cells and imaged at specific time points during induction (Figure 5B-D); and second, we induced cells and used time-lapse imaging to monitor GFP-Tub3 production and dynamics in living cells (Video S1). These experiments show that the total cellular GFP-Tub3 signal increases from two to three hours of induction, but is slightly decreased at 24 hours of induction (Figure 5C). The accumulation of GFP-Tub3 into microtubules shows a different trend. GFP-Tub3 is detectable in microtubule polymer approximately an hour after galactose induction, before it is detectably increased in the cytoplasm, and the amount of GFP signal in microtubules steadily increases to reach the highest level 24 hours after induction (Figure 5B, C). In addition to the accumulation of GFP-Tub3 in microtubules, we observed the formation of GFP-Tub3 assemblies that are separate from microtubules (Figure 5B). These foci are reminiscent of the tubulin assemblies observed during Tub1/α-tubulin overexpression in Figure 4D. Interestingly, GFP-Tub3- containing assemblies are not detectable by 24 hours post induction (Figure 5B). Together these results suggest that cells may limit the accumulation of α-tubulin protein during overexpression, and identify α-tubulin-containing assemblies as a potential intermediate state involved in regulating α-tubulin levels.

**Figure 5:**
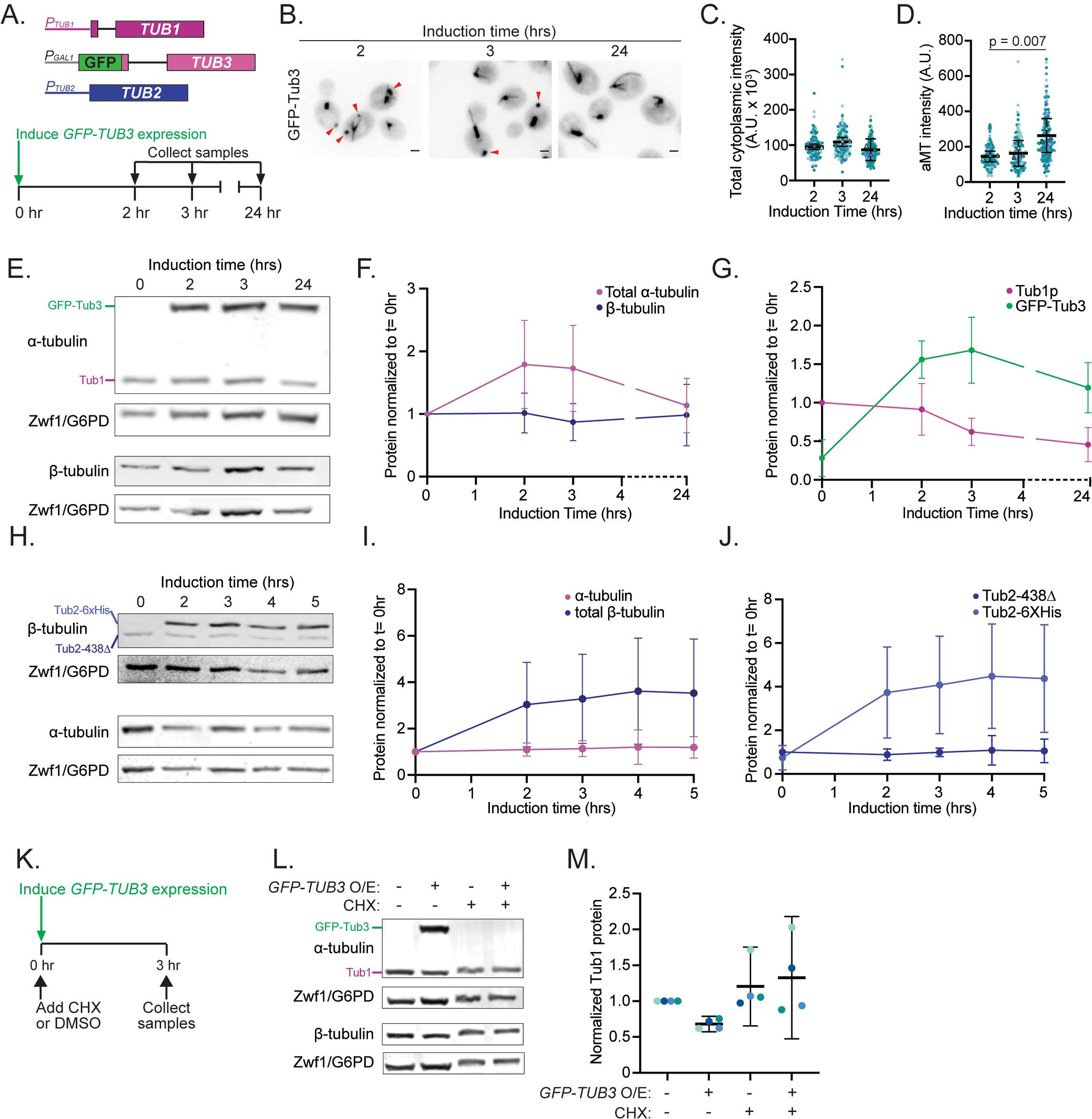
Tub1 and Tub3 isotypes are balanced to set α-tubulin protein levels A. Schematic of gene composition for the GFP-Tub3 induction strain and time course of induction experiments. B. Example images of GFP-Tub3 in cells at indicated induction time. Arrowheads indicate tubulin assemblies. Scale bar = 1 µm. Images are a maximum intensity projection of z series. C. Quantification of whole cell GFP-Tub3 fluorescence intensity. Each dot represents a single cell and are colored by replicate, triangles represent the mean of each experiment. Bars are mean ± 95% CI. D. Quantification of GFP-Tub3 fluorescence intensity in astral microtubules. Each dot represents a single microtubule in a single cell and are colored by replicate, triangles represent the mean of each experiment. P-value between wild type and mutant based on t-test after one-way ANOVA. Bars are mean ± 95% CI. E. Representative western blot of GFP-Tub3 induction. Cells were blotted for α- or β-tubulin and Zwf1 (G6PD) as a loading control. F. Quantification of total α- and β-tubulin levels across GFP-Tub3 induction. Dots are mean ± SD from five independent experiments, normalized to the level of α- or β-tubulin prior to induction. G. Quantification of Tub1p and GFP-Tub3 levels across induction. Dots are mean ± SD from five independent experiments, normalized to the Tub1p levels prior to induction. H. Representative western blot of Tub2-6xHis induction. Cells were blotted for α- or β- tubulin and Zwf1 (G6PD) as a loading control. I. Quantification of total α- and β-tubulin levels across Tub2-6xHis induction. Dots are mean ± SD from five independent experiments, normalized to α- or β-tubulin levels prior to induction. J. Quantification of Tub2-438Δ and Tub2-6xHis levels across Tub2-6xHis induction. Dots are mean ± SD from five independent experiments, normalized to tub2-438Δ levels prior to induction. K. Scheme for GFP-Tub3 induction and cycloheximide treatment. L. Representative western blot of GFP-Tub3 induction with or without cycloheximide (CHX) treatment. M. Quantification of Tub1 protein levels in indicated treatment groups. Data represent four experiments and are colored by day. Bars are mean ± 95% CI.

Our time-lapse imaging captures the formation and dissolution of the GFP-Tub3- containing assemblies. We find that puncta of GFP-Tub3 signal begin to appear around two hours post induction and diffuse around the cell before dissolving at around 5.5 hours post induction (Video S1-3). We note that the signal of GFP-Tub3 in microtubules gradually increases over this time course. These results indicate that GFP-Tub3-containing assemblies may represent transient stores of α-tubulin en route to heterodimer formation and/or microtubule polymerization.

We next compared how the induction of high levels of GFP-Tub3 expression impacts the levels of the other α-tubulin isotype, Tub1. Our western blots show that total α-tubulin is increased approximately two-fold at 2 hours post induction, followed by a gradual decrease that returns to pre-induction levels by 24 hours (Figure 5E, F). When we compare individual levels of the GFP-Tub3 and Tub1 isotypes by western blot, we see that as GFP-Tub3 levels strongly increase during the first two hours after induction, levels of endogenous Tub1 decrease (Figure 5G). After 24 hours of induction, GFP-Tub3 levels decreased to nearly match the level of Tub1 that we measured prior to galactose-induction, while Tub1 was decreased to less than 0.5x of pre-induction levels (Figure 5F). Through this time course we find no change in β-tubulin levels, even by 24 hours (Figure 5E, F). The consistent level of β-tubulin stands in contrast to the loss of Tub1 and indicates that loss of Tub1 is not attributable to titration of protein levels through cell division. These data suggest that when expression of one α-tubulin isotype is strongly increased the levels of the other α-tubulin isotype decreases in response, and as a result the total α-tubulin is transiently elevated before returning to pre-induction levels.

We next asked if β-tubulin protein levels are similarly balanced in response to increased β-tubulin expression. To answer this question, we created a strain where we could distinguish the endogenous β-tubulin from conditionally overexpressed β-tubulin by western blot. We utilized a haploid strain in which the last 19 codons of the carboxy-terminal tail were removed from chromosomal *TUB2*, creating a *tub2*-438Δ allele, and transformed these cells with a plasmid for galactose-inducible expression of full-length Tub2 with a 6xHis tag fused to the carboxy-terminus. This allows us to distinguish the faster-migrating, endogenous tub2-438Δ polypeptide from the slower-migrating, exogenous Tub2-6xHis on a western blot (Figure 5H). Due to the toxicity associated with excess β-tubulin expression, we used an abbreviated time course lasting 5 hours for our western blots. Nevertheless, we see a steady increase in total β- tubulin levels during galactose induction, reaching a four-fold increase by 4 hours (Figure 5I). The induced Tub2-6xHis increases steadily during this time course, while the endogenous tub2- 438Δ β-tubulin shows no change in protein levels (Figure 5J). We also see no change in the α- tubulin levels over the time course of Tub2-6xHis expression (Figure 5I). This suggests that β- tubulin expression does not equilibrate like α-tubulin expression.

Finally, we wanted to determine if the coordination of α-tubulin levels between the Tub1 and Tub3 isotypes was stimulated by changes in mRNA levels or protein levels. To test this, we induced GFP-Tub3 expression for three hours in cells treated with cycloheximide to inhibit translation or DMSO as a control, and took samples for western blotting (Figure 5K, L). We predicted that if increased GFP-*TUB3* mRNA levels are sufficient to trigger the decline in Tub1 protein levels, then inducing GFP-*TUB3* transcription in the presence of cycloheximide would be sufficient to trigger the depletion of Tub1. We instead found that cells treated with cycloheximide and induced for GFP-Tub3 expression maintained steady Tub1 levels (Figure 5M). Together these data suggest that cells regulate α-tubulin protein levels by maintaining an equilibrium between α-tubulin proteins. When protein levels of a new α-tubulin isotype increase, the other α- tubulin isotype is gradually lost from the cell until total α-tubulin returns to a predetermined set point.

## Discussion

All eukaryotic genomes contain families of genes for α- and β-tubulins that typically exhibit developmental and cell-type specific expression programs (Erickson 2007; Wickstead and Gull 2011; Findeisen et al. 2014; F. D. Miller et al. 1987). This raises a fundamental question of how the expression of these genes and proteostasis of α- and β-tubulins may be coordinated to meet a cell’s demand for tubulin heterodimers, while maintaining the appropriate balance of each subunit. In this study we identify an asymmetric requirement for tubulin gene copy number in budding yeast; cells have a stronger requirement for α- over β-tubulin genes.

We find that the soluble tubulin pool contains ∼25% more α- tubulin than β-tubulin and there are distinct consequences when the normal ratio of α- to β-tubulin is disrupted. Increased β-tubulin levels result in a loss of microtubules and formation of non-microtubule tubulin assemblies.

Conversely, increased α-tubulin is tolerated through an equilibration mechanism that decreases levels of an alternative α-tubulin isotype to return to a predetermined level of total α-tubulin. Our findings indicate that α- and β-tubulin are regulated through distinct mechanisms and identify an important role for α-tubulin in preventing β-tubulin toxicity.

In principle, α- and β-tubulin proteins exist in multiple states in the cell, including αβ- tubulin heterodimers that are in equilibrium between a soluble pool and microtubule polymer, and non-heterodimer states that represent steps in tubulin biogenesis or recycling and could be present as soluble monomers (Figure 6). Decreasing α- or β-tubulin gene copy number would be expected to undersupply the corresponding tubulin protein and deplete one or more of these states, manifesting as less tubulin detected in the soluble pool and/or short, slowly polymerizing microtubules. Our results show that decreasing gene copy number does limit microtubule length and slow polymerization, with β-tubulin genes having a stronger impact than α-tubulin genes (Figure 2). This suggests that in yeast β-tubulin is limiting for the heterodimer state. However, loss of α- or β-tubulin genes has little effect on the amount of corresponding tubulin detected in the soluble pool (Figure 2). We estimate that diploid cells maintain ∼0.31 µM α-tubulin and ∼0.25 µM β-tubulin in the soluble pool, based on an average volume of 120 fL for a diploid budding yeast cell (Figure 2G; Sherman, 2002). Overall, this supports a model in which cells maintain a relatively constant concentration of α- and β-tubulin in the soluble pool, and undersupplying tubulin by decreasing α- or β-tubulin gene copy number primarily affects the microtubule polymers.

**Figure 6:**
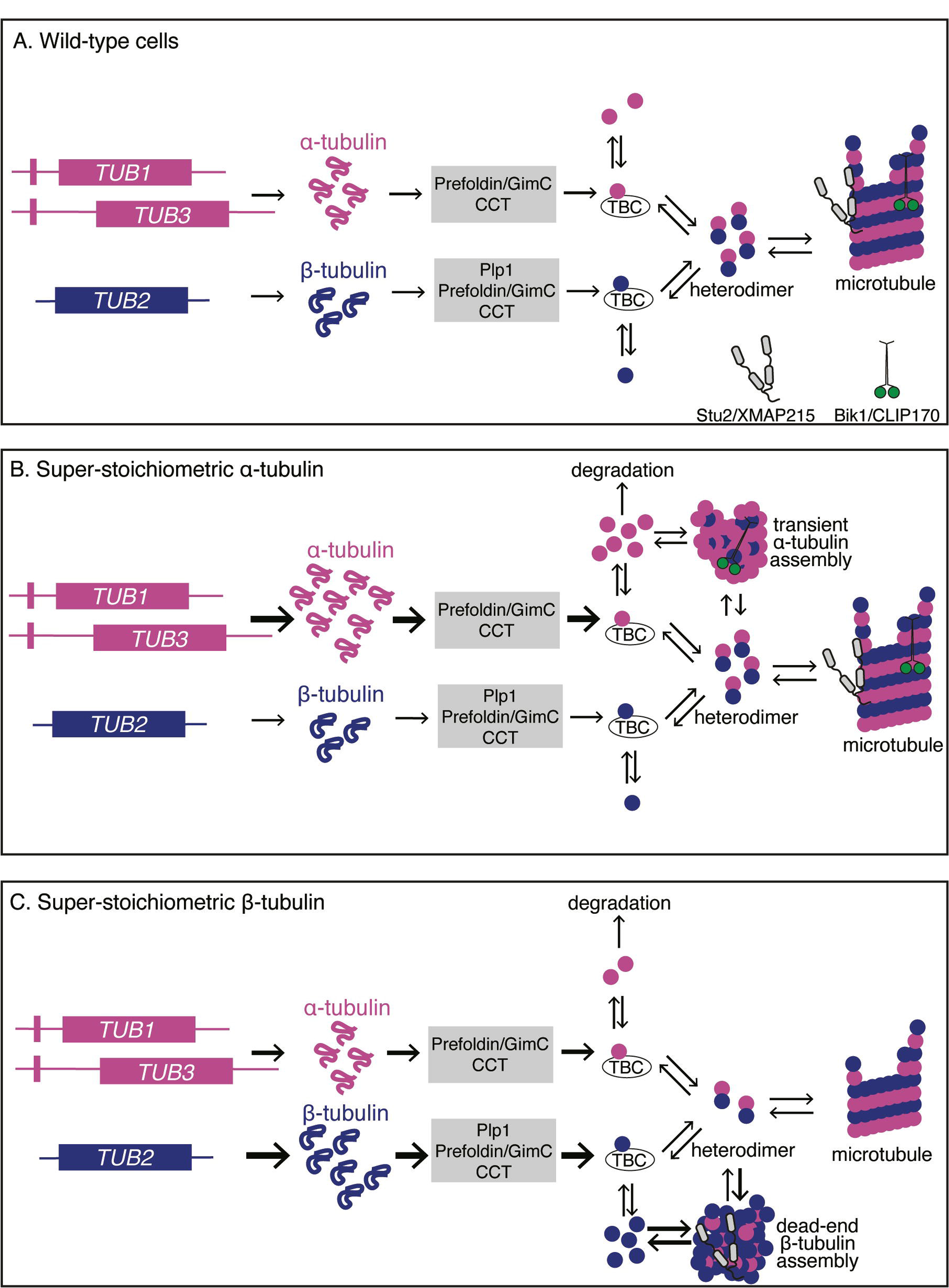
Proposed model for α-tubulin regulation of β-tubulin A. Updated model for how wild-type cells express tubulin from mRNA to protein and undergo folding and assembly into heterodimers. Our data suggests that cells hold an excess of α-tubulin that could exist as TBC-bound or soluble monomers. B. In conditions where there is super-stoichiometric α-tubulin in cells, surplus α-tubulin drives monomerization and the formation of a transient α-tubulin assemblies. Our data shows that cells then down regulate the alternative isotype to restore endogenous expression and at the same time the assemblies dissipate. C. In conditions where there is super-stoichiometric β-tubulin in cells, surplus β-tubulin drives the monomerization and formation of dead-end β-tubulin assemblies. Our data suggests these assemblies disrupt microtubule architecture and +TIP localization.

It is important to note that our measurements of tubulin in soluble lysates is likely an underestimate of the total number of tubulin molecules in the cell. Previously published tomograms of anaphase spindles in diploid *S. cerevisiae* show a total of ∼23 µm of microtubule polymer in the spindle (Winey et al. 1995). Assuming 1625 heterodimers per micrometer of a 13-protofilament budding yeast microtubule lattice (Howes et al. 2018), we estimate that the amount of microtubule lattice in these anaphase spindles would require at least 37,375 heterodimers in the polymer state. Our measurements of soluble tubulin after cell lysis indicate an average of 23,242 α-tubulins and 18,959 β-tubulins; therefore, our experiments either underrepresent anaphase cells or our soluble fractions may not include all of the tubulin in spindle microtubules (Figure 2G). We do suspect that some tubulin in microtubule polymers is lost to the insoluble fraction after lysis, because we find in separate experiments that chilling cells at 4°C for one hour to completely depolymerize microtubules prior to lysis leads to an increased but highly variable amount of α- and β-tubulin in the soluble fraction (L. Wethekam, unpublished observation). Thus, we regard our lysate measurements as an accurate reflection of the soluble tubulin in cells, rather than the total tubulin.

As further evidence of cells maintaining soluble pool at the expense of microtubule polymers, we find that pre-anaphase spindles are shorter in cells lacking a copy of α- or β- tubulin encoding genes, individually or in combination (Figure 1E, F). These shorter spindles are consistent with a decrease in microtubule polymer in the cell – we noted correlative changes in astral microtubule length and pre-anaphase spindle length across genotypes, suggesting an equilibrium between the cytoplasmic and nuclear pools of tubulin heterodimers that is equally affected by changes in gene copy number. Despite the effect of α- or β-tubulin gene loss on microtubule polymers, our fitness assay shows that this effect alone does not strongly impact fitness (Figure 1B, C). This suggests that budding yeast cells produce more tubulin heterodimers than necessary to build a functional spindle, at least under optimal culturing conditions. In other words, supply exceeds demand and yeast cells maintain a surplus of α- and β-tubulin.

How do cells set the level of tubulin surplus? Mammals possess a defined autoregulatory mechanism for β-tubulin that responds to increases in soluble tubulin and prevents additional tubulin production through targeting tubulin mRNA for decay via the ribosome-associated protein TTC5 (Bachurski et al. 1994; Cleveland and Havercroft 1983; Lin et al. 2020). Budding yeast lack a clear homologue for TTC5, calling into question whether autoregulation is broadly conserved across species. Our results suggest that budding yeast limit α-tubulin accumulation through an alternative mechanism. Constitutively supplying cells with an extra copy of *TUB1* does not increase soluble tubulin levels or strongly affect microtubule assembly (Figure S3).

Conditionally inducing high levels of transcription of the *TUB3* α-tubulin isotype stimulates a decrease in protein levels of the alternative α-tubulin isotype Tub1, and this requires the translation of the *TUB3* mRNA (Figure 5). We find that these cells eventually return to total levels of α-tubulin that are similar to basal levels. Based on these results, we propose a model for α-tubulin regulation that is different from autoregulation. As depicted in Figure 6, we propose that when α-tubulin is overexpressed, the surplus α-tubulin exists as tubulin monomers and assemblies that can either exchange with αβ-heterodimers or be targeted for degradation (Figure 6). In this way, a newly created α-tubulin would exchange with any pre-existing α-tubulin in the cell and promote its degradation. The tubulin assemblies that we observe upon α-tubulin overexpression may represent a transient enrichment of the excess α-tubulin, which is subsequently incorporated into microtubules or degraded. We speculate that exchange with the excess state of α-tubulin may be an important part of tubulin proteostasis and could exist as either monomers or bound to TBCs that allow for the recycling of αβ-heterodimers and preventing the accumulation of monomeric β-tubulin.

Our results indicate that budding yeast regulate β-tubulin differently than α-tubulin. β- tubulin does not appear to access an alternative state that can be rapidly turned over. When we conditionally induce *TUB2* transcription, soluble β-tubulin levels increase without a corresponding decline in the levels of the native β-tubulin or an increase in microtubule polymer (Figure 5J). The autoregulation model predicts that this increase in soluble tubulin should induce mRNA decay and thereby inhibit further translation, creating a steady state level of β-tubulin. In our experiments, β-tubulin levels continue to increase for hours without reaching steady state levels (Figure 3E-M). In contrast to α-tubulin, we propose that super-stoichiometric β-tubulin forms ‘dead-end’ assemblies that exhibit very slow exchange and are not readily targeted for degradation (Figure 6). Furthermore, super-stoichiometric β-tubulin may co-assemble with αβ- heterodimers to form highly stable, non-microtubule assemblies, which could explain why even a small excess of β-tubulin is sufficient to deplete microtubules. Together, this evidence suggests that budding yeast are unlikely to use the autoregulatory mechanism found in metazoans. Whereas autoregulation may have evolved in metazoans to manage the expanded isotype repertoire and programmed changes in tubulin expression, budding yeast lack autoregulation are therefore exquisitely sensitive to increases in β-tubulin expression.

While overexpression represents an extreme example of imbalanced α- and β-tubulin, our experiments in heterozygous knockout mutants demonstrate toxicity linked to a modest increase in β-tubulin. Removing a single copy of *TUB1* creates an 36% increase in β-tubulin in the soluble pool (Figure 2G-H). This increase could represent the emergence of monomeric β- tubulin after saturating its binding partners, α-tubulin and TBCA/Rbl2. How might monomeric β- tubulin be toxic? We speculate that the tubulin assemblies formed when β-tubulin is in excess could involve interactions between β-tubulin monomers and/or between β-tubulin monomers and αβ-heterodimers. In this scenario, any β-β contacts along the longitudinal interface would lack the catalytic residues that α-tubulin normally provides to the exchangeable GTP-binding site (Anders and Botstein 2001; LaFrance et al. 2022), and therefore these assemblies would be locked in a high affinity GTP-bound state. These hyperstable assemblies could then sequester tubulin heterodimers and select MAPs such as XMAP215/Stu2, depleting components from the cell’s microtubule networks. Our conditional overexpression experiments suggest that even a small excess of monomeric β-tubulin may be sufficient to set off this cascade and lead to microtubule loss.

The asymmetric requirement for α- vs β-tubulin genes is also seen in metazoans. As multicellular species evolved, so did the requirement for robust tubulin expression. Some of the clearest examples of this come from studies of brain development in mouse models, where differentiating cells experience an increased demand for αβ-tubulin. Knocking out the α-tubulin gene *TUBA1A* causes severe brain malformation and is perinatal lethal, but knocking out any of several β-tubulin genes results in milder malformations or subsequent axon repair defects (Bittermann et al. 2019; Latremoliere et al. 2018). Missense mutations in any of these tubulin isotypes are linked to severe brain malformations in humans, demonstrating the important roles of these genes in supplying functional tubulin (Bahi-Buisson and Cavallin 1993; Romaniello et al. 2018; Park et al. 2021). That the full knockouts of β-tubulin genes are less severe than knocking out α-tubulin *TUBA1A* indicates a stronger requirement for α-tubulin gene copy number than β-tubulin gene copy number, reminiscent of our findings in the single celled yeast. How could cells ensure sufficient α-tubulin expression to buffer against β-tubulin induced toxicity? One way is through simply having more genes for α-tubulin than β-tubulin. Indeed, we find that in most species the number of α-tubulin genes is greater than or equal to the number of β-tubulin genes, suggesting unbalanced β-tubulin gene expansion is rare and perhaps detrimental (Findeisen et al. 2014). In *S. cerevisiae* the greater number of α-tubulin genes arose from a whole genome duplication that resulted in two α-tubulin isotypes: *TUB1* and *TUB3*. While both α-tubulins were maintained, the duplicate β-tubulin was lost from the genome leaving only *TUB2*. Additional copies of *TUB2* or aneuploidies of its chromosome (chromosome VI) are lethal, suggesting that loss of the second β-tubulin gene millions of years ago may have been advantageous (Katz, Weinstein, and Solomon 1990; Burke, Gasdaska, and Hartwell 1989; Torres et al. 2007; Anders et al. 2009). Further back in the evolution of tubulin, we know that α- and β-tubulin are themselves the result of a gene duplication and diversification of the bacterial protein FtsZ. Perhaps an important step in this evolution from a monomer to a heterodimer was the emergence of an α-tubulin precursor to regulate the activity and prevent the toxicity of the β- tubulin precursor.

## Methods

### Yeast manipulation and culturing

Yeast manipulation, media, and transformations were performed by standard methods (Amberg, Burke, and Strathern 2000). A detailed list of strains and plasmids is provided in Supplementary Tables 1 and 2. Deletion mutants were generated by PCR-based, homologous recombination methods (Petracek and Longtine 2002). Spc110-mNeonGreen, Spc110-tdTomato, and Kip3- mNeonGreen were generated using PCR-based methods and expressed from the genomic locus (Sheff and Thorn 2004). The mNeonGreen fluor was provided by Allele Biotechnology and Pharmaceuticals (San Diego, CA) (Shaner et al., 2013). Bik1-3GFP, Bim1-3GFP, and Stu2- 3GFP were generated using an integrating plasmid and expressed from the genomic locus. The Bik1-3GFP integrating plasmid was a gift from Dr. David Pellman (Harvard University; Carvalho *et al*., 2004). The Bim1-3GFP integrating plasmid was a gift from Dr. Tim Huffaker (Cornell University; Wolyniak *et al*., 2006). The Stu2-3GFP integrating plasmid was generated by PCR-amplification of base pairs 1479-2664 from chromosomal STU2 and cloning into the BamHI site of pBJ1376 (Lee, Oberle, and Cooper 2003), to create plasmid pJM459. This plasmid was digested with EcoRV for integration at the *STU2* locus and confirmed by PCR. GFP-Tub1 fusions were integrated using plasmid pSK1050 (Song and Lee 2001) and expressed in addition to the native α-tubulin. To build the inducible Tub2 expression plasmid, we first built a diploid strain with one allele of wild-type Tub2 and the other allele tagged with 6xHis. This strain was generated by transforming a PCR fragment containing the c-terminal 6xHis tag with 331 base pairs of 3’ UTR followed by a TRP1 marker (pJM385, Li and Moore, 2020). We confirmed that this Tub2-6xHis allele rescues β-tubulin function in yeast. Subsequently, a PCR fragment from pFA6a-KanMX6-PGAL1 containing homology to replace 185 base pairs 5’ of the start codon was transformed into the *tub2-6xHis*/*TUB2* strain. A strain with both the promoter and the 6xHis tag on the same allele was used a template. Finally, to construct the plasmid, a fragment of DNA containing the galactose promoter, the *TUB2* coding sequence, and 427 base pairs of the 3’ UTR (including the TRP1 marker) were cloned into the NotI and SacI sites of pRS316 or pRS313. To build the inducible Tub1 plasmid, a diploid strain was generated that replaced 166 base pairs of the endogenous promoter of one allele of *TUB1* with KanMX6-PGAL1. To assemble the plasmid a fragment of DNA containing the galactose promoter, the *TUB1* sequence (including intron), and 487 base pairs of the 3’ UTR were cloned into the NotI site of pRS315 or the NotI and SacI sites of pRS316. To build the extra copy of *TUB1* a fragment of 992 base pairs 5’ of *TUB1* through 487 base pairs of the 3’ UTR including the intron were cloned into the NotI and KpnI sites of pRS316.

### Microscopy

All live-cell fluorescence images were collected by spinning disk confocal microscopy using a Nikon Ti-E microscope equipped with a 1.45 NA 100x CFI Plan Apo objective, piezo electric stage (Physick Instrumente, Auburn, MA), spinning disk confocal scanner unit (CSU10: Yokogawa), 488 and 561 nm lasers (Agilent Technologies, Santa Clara, CA), and an EMCCD camera (iXon Ultra 897, Andor Technology, Belfast, UK); using NIS Elements software (Nikon, Tokyo, Japan). All fixed-cell images and time-lapse DIC images were collected on a Nikon Ti-E wide field microscope equipped with a 1.49 NA 100xCFI160 Apochromat objective and an ORCA-Flash 4.0 LLT sCMOS camera (Hammamatsu Photonics, Japan) using NIS Elements software (Nikon, Tokyo, Japan). For live cell imaging on both microscopes, stages were incubated at 30°C using an ASI 400 Air Stream Incubator (NEVTEK, Williamsville, VA).

### Liquid Growth Assay

Cells were grown in 3 mL of rich liquid media (YPD) to saturation at 30°C and diluted 50-fold into fresh media. The diluted cultures were then aliquoted into a 96-well plate, with three to six technical replicates per experiment, and incubated at 30°C with single orbital shaking in a Cytation3 plate reader (BioTek, Winooski, VT). The OD600 was measured every 5 minutes for 24 hours. Doubling time was calculated by fitting the growth curves to a nonlinear exponential growth curve as previously published (Fees and Moore, 2018; MatLab R2018a; Mathworks, Natick, MA). Doubling times were normalized to the mean of all technical replicates from wild-type controls. P-values are from student’s t-test after a one-way ANOVA with a Tukey post-hoc test for p < 0.05, or the Tukey post-hoc test for p >0.05.

### Solid Growth Assays

Cells were grown in rich liquid media to saturation at 30°C, and a 10-fold dilution series of each culture was spotted to either rich media plates or rich media plates supplemented with 5 or 10 µg/mL benomyl (Sigma Aldrich #381586, St. Louis, MO). Plates were grown at the indicated temperature for the indicated days. For the tubulin induction experiments, cells were grown in dropout media to select for the expression plasmid to saturation at 30°C, and a 10-fold dilution series of each culture was spotted to drop out plates supplemented with either 2% glucose to inhibit induction or 2% galactose to induce expression.

### Pre-anaphase spindle measurements

Cells expressing Spc110-mNeonGreen were grown in rich liquid media at 30°C, then diluted and grown to log phase in fresh media. Cell were adhered to coverslips coated with concanavalin A and left in nonfluorescent media for imaging (Fees, Estrem, and Moore 2017). Cells were imaged at 30°C. Z-series consisting of a 6 µm range at 0.35 µm steps were acquired every 20 seconds for 5 minutes.

Pre-anaphase cells were identified in image series of asynchronous cultures based on bud size and spindle length, and cropped. Z-series images were processed in FIJI (Wayne Rasband, NIH) using the “Despeckle” plugin followed by the “Remove Outlier” plugin with the radius set to 1 pixel and the threshold 25 to remove bright, outlier pixels. Spindle length over time was determined using a custom analysis program (Thomas, Ismael, and Moore 2020). This program determines the X,Y,Z coordinates of the brightest pixel within the three-dimensional image stack (i.e., the first SPB), applying a Gaussian blur around this pixel, then identifying the second brightest pixel in the image stack (the second SPB). Spindle length was then defined as the linear distance between these points in three dimensions. This process was then repeated for every Z-series at each point in the time course. P-values are from student’s t-test after a one-way ANOVA with a Tukey post-hoc test for p < 0.05.

### Microtubule dynamics in living cells

Cells expressing Bik1-3GFP were grown in rich liquid media or selective drop out media at 30°C, then diluted and grown to log phase in fresh media. Cells were adhered to coverslips coated with concanavalin A and left in nonfluorescent media for imaging. Cells were imaged at 30°C at 5 second intervals for 10 minutes. Z-series consisting of a 7 µm (2A-D) or 6 µm (Supplement 1D,E) range with a step size of 0.45 µm was taken at each timepoint and analyzed as a 2-D maximum intensity projection in FIJI. Lengths were measured from the edge of the SPB to the tip of the astral microtubule at each time point. All analyses were done in asynchronous pre-anaphase cells. Lengths in pixels were processed using a custom analysis program (Estrem, Fees, and Moore 2017). P-values are from student’s t-test after a one-way ANOVA with a Tukey post-hoc test for p < 0.05.

### Immunofluorescence

Cells with inducible *TUB1* or *TUB2* expression plasmids were grown in selective drop out media supplemented with 2% raffinose overnight. Cells were diluted into fresh medium, grown to log phase, and then induced with the addition of 2% galactose. The fixation and immunostaining protocol is modified from a previously published method (R. K. Miller 2004). Cells were fixed with 3.7% formaldehyde (Sigma-Aldrich, 252549) at 30°C for 2 hours, centrifuged at 1,400xg for 3 minutes, washed twice with wash buffer (40 mM KPO4, pH 6.5), and then stored overnight at 4°C. The fixed cells were then washed twice with wash buffer plus 1.2 M sorbitol and digested with 10 µl of 20T 50 mg/ml zymolyase (Nacali Tesque, 07663-91) supplemented with 15 µl β-mercaptoethanol (Sigma-Aldrich, M3148) for 45 minutes at 37°C. Digested cells were spun down at 600xg for 3 minutes, washed once with wash buffer plus 1.2 M sorbitol, and then resuspended in 20µl wash buffer plus 1.2M sorbitol. 20µl of cell suspension was spotted onto each well of a Teflon coated 10-well slide (Polysciences, 18357) that had been pre-treated with 10 ng/µl poly-L-lysine. Cells adhered for 10 minutes at room temperature. Liquid was aspirated off before immediately permeabilizing the cells in a coplin jar of cold methanol (Acros Organics, 444310050) for 6 minutes, followed by immersing the slide in cold acetone (Fisher Chemical, A18-4) for 30 seconds. Cells were blocked at room temperature for 1 hour in blocking buffer (1X PBS + 0.5% BSA), then incubated overnight at 4°C in a humid chamber with mouse anti-α-tubulin (4A1; 1:100 in blocking buffer) or mouse anti-β-tubulin (E7, undiluted culture media collected from the hybridoma cells). Wells were washed 4 times for 10 minutes with blocking buffer, then incubated at room temperature for 1 hour in a dark humid chamber with goat anti-mouse IgG-Alexa488 (Invitrogen, A11001, 1:500 in blocking buffer). Cells were washed 4 more times with blocking buffer. DAPI mounting solution (Vector Laboratories, H-1200) was added to the slide. Slides were imaged on the widefield microscope described above. Z-series consisting of a 7 µm range at 0.5 µm steps were acquired for Alexa488 and DAPI, with DIC image at home position. For analysis, in-focus planes were identified and maximum intensity projections were created.

### Cold Sensitivity Assay

Cells expressing GFP-Tub1 Spc110-tdTomato and the pGal-Tub2-6xHis plasmid were grown in 3 mL of selective growth media supplemented with 2% raffinose at 30°C, diluted into fresh medium, and grown to mid-log phase. Cells were then induced with the addition of 2% galactose for 3 hours at 30°C. As a control a second set of cultures were supplemented with 2% glucose to block induction. Cells were then moved to 4°C at 0, .25, .5, 1, 2, and 24 hours of induction and then fixed with 3.7% formaldehyde in 0.1 M KPO4 for two minutes. The cells were pelleted in a benchtop mini microcentrifuge, resuspended in a quencher solution (0.1% Triton-X, 0.1 M KPO4, and 10 mM ethanolamine, pelleted again, and washed once in 0.1 M KPO4. Fixed cells were loaded into slide chambers coated with concanavalin A, washed with 0.1 M KPO4, and the chambers were sealed with VALAP. Z-series were acquired using a 7 µm range separated by µm steps on the widefield microscope described above. For analysis, maximum intensity projections were created to score astral microtubules and tubulin assemblies.

### +TIP behavior assay

Cells expressing the indicated GFP- or mNeonGreen-tagged +TIP with mRuby-Tub1 and *TUB1* or *TUB2* induction plasmids were grown overnight in drop out growth media supplemented with 2% raffinose at 30°C, then diluted back into fresh medium and grown to log phase. Cells were induced with 2% galactose, or control cells were not induced. At 2 hours of induction cells were harvested and adhered to coverslips coated with concanavalin A. Cells were imaged 30°C at 10 second intervals for 1 minute with a 5.4 µm range with at 0.45 µm step size. For analysis, 2-D maximum intensity projections were generated in FIJI. Cells were scored for ‘+TIP activity’, which was defined by the presence of a focus of labeled +TIP that tracked the plus end of a microtubule as it grew and/or shortened.

### Western blotting

To prepare soluble protein lysate, log-phase cells were pelleted and resuspended in 2 M Lithium acetate and incubated for 5 minutes at room temperature. Cells were pelleted again and resuspended in 0.4 M NaOH for 5 minutes on ice. Cells were pelleted and resuspended in 2.5x Laemmli buffer and boiled for 5 minutes. Before loading gels, samples were boiled and centrifuged at 6,000xg for 3 minutes. Before blotting (for all westerns in Figures 3-5) total protein concentration of clarified lysate was determined by Pierce 660 nm protein assay with the Ionic Detergent Compatibility Reagent (Cat. 1861426 and 22663, Rockford, IL). ∼10 ug of total protein were then loaded. Samples were run on 10% Bis-Tris PAGE gels in NuPAGE MOPS running buffer (50 mM MOPS, 50 mM TrisBase, 0.1% SDS, 1 mM EDTA, pH 7.7) at 0.04 mAmp per gel for 1 hour, or 1.5 hours to separate β-tubulins in Figure 5H-J. Gels were transferred to PVDF (Millipore, IPFL85R) in NuPAGE transfer buffer (25 mM Bicine, 25 mM Bis-Tris, 1 mM EDTA, pH7.2) at 0.33 mAmp for 1 hour. Membranes were then blocked for 1 hour at room temperature in PBS blocking buffer (LI-COR, 927-70001). Membranes were probed in PBS blocking buffer including the following primary antibodies: mouse-anti-α-tubulin (4A1; at 1:100; Piperno and Fuller, 1985), mouse-anti-β-tubulin (E7; at 1:100; Developmental Studies Hybridoma Bank, University of Iowa), rabbi-anti-Zwf1 (Glucose-6-phosphate dehydrogenase; Sigma A9521; at 1:10,000) overnight at 4°C. After incubation in primary antibody, membranes were washed once in PBS for 5 minutes at room temperature and then probed with the following secondary antibodies: goat-anti-mouse-680 (LI-COR 926-68070, Superior, NE; at 1:15,000) and goat-anti-rabbit-800 (LI-COR 926-32211; at 1:15,000) for 1 hour at room temperature. After incubation in secondary antibodies, blots were washed twice in PBST (1XPBS, 0.1% Tween- 20), once in PBS, and imaged on an Odyssey Imager (LI-COR, 2471).

### Quantifying tubulin concentration

To determine levels of soluble tubulin, wild-type or heterozygous null cells were grown to log phase in rich media at 30°C. To prepare lysate of 5x10^7^ cells in 50 µL samples, cells were counted on a hemocytometer and the appropriate volume of cells was harvested and prepared as described above. Lysate was resuspended in 2.5x Laemmli buffer and standards of purified yeast tubulin were prepared by diluting protein to 2.5 ng/µL in 2.5x Laemmli buffer. Samples containing increasing amounts of cells (3.5, 4.5, 6, and 8 x 10^6^) and purified protein (4,10, 15, 30 and 40 ng) were loaded and blotted as described above. Band intensities were quantified using the gel analysis plug-in in FIJI, and signal intensities were converted to nanograms using the standard curve, and converted to molecules/cell with the following:

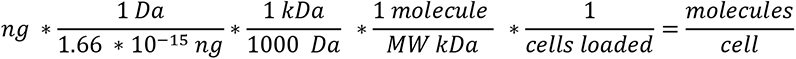

Each of the molecule/cell values for a biological replicate were averaged for a single experiment with the corresponding α- and β-tubulin values paired. P-values are from student’s t-test after a one-way ANOVA with a Tukey post-hoc test for p < 0.05.

### Colony Formation Assay

Cells were grown overnight in drop out growth media supplemented with 2% raffinose. Cells were then diluted in fresh media containing 2% raffinose, grown to log phase, and induced by the addition of 2% galactose to the culture. At the indicated time points, samples were collected from the cultures and cell density was counted on a hemocytometer. Based on these counts, cells dilutions were adjusted in order to plate the same number of expected cells for each time point and condition. For the empty vector, *TUB1* overexpression, and *TUB1* and *TUB2* simultaneous overexpression conditions, 300 cells were spread on each plate. For *TUB2* overexpression alone, 500 cells were spread on each plate to enhance the sensitivity of the assay.

### pGAL-GFP-Tub3 induction timepoint imaging

Cells containing *pGAL1-GFP-TUB3* at the endogenous *TUB3* locus were grown in rich media supplemented with 2% raffinose at 30°C, diluted back into fresh medium and grown to log phase, and then induced with 2% galactose. For single timepoint imaging: at 2, 3 and 24 hours post induction, cells were harvested, suspended in nonfluorescent media and adhered to coverslips coated with concanavalin A. Z-series images consisting of 8 µm range in 0.45 µm steps were collected. Images were analyzed in FIJI. For time-lapse imaging: after 40 minutes of induction cultures were harvested and adhered to slides coated with concanavalin A and left in nonfluorescent media containing 2% galactose for imaging. Z-series images consisting of 6 µm range in 0.45 µm steps were captured at two minute intervals for six hours. P-values are from student’s t-test after a one-way ANOVA with a Tukey post-hoc test for p < 0.05.

## Supplemental Material

Supplemental materials include five figures, three videos related to Figure 5, a yeast strain table, a table of plasmids used in the study, and a table of oligonucleotides used in the study.

## Acknowledgments

We are grateful to members of the Moore lab for helpful advice and discussions. This work was supported by NIH R01 GM112893, R35 GM 136253 (J.K.M.) and L.W. was supported by T32 GM136444 and the Bolie Scholar Award.

## Author contributions

Linnea C. Wethekam: Conceptualization, formal analysis, investigation, methodology, validation, visualization, writing – original draft, writing – review & editing Jeffrey K. Moore: Conceptualization, funding acquisition, methodology, project administration, resources, supervision, visualization, writing – original draft, writing – review & editing

---

**Supplemental Figure 1:**
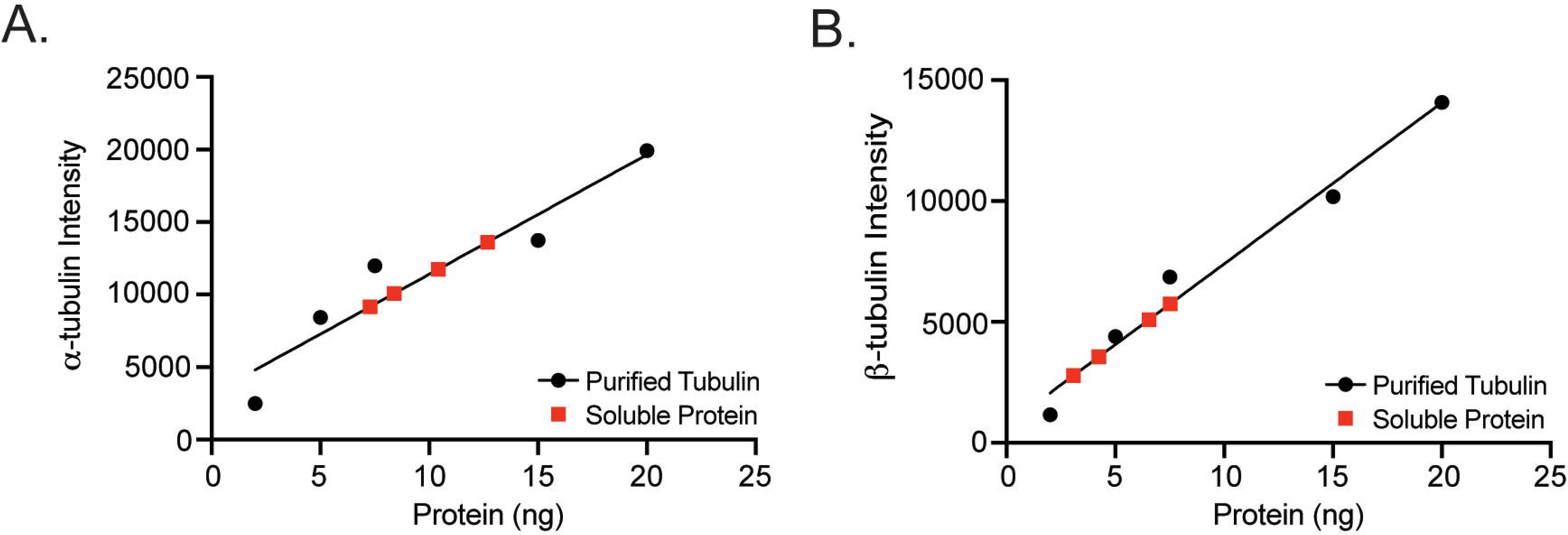
Example standard curves for determining molecules of α- or β-tubulin per cell. A. Example standard curve of α-tubulin intensity vs amount of purified protein loaded. Black line is the linear best fit. Red squares are the intensity measurements from 3.5, 4.5, 6, and 8 x10^6^ number of cells, fit to the standard curve. B. Example standard curve of β-tubulin intensity vs amount of protein loaded. Black line is the linear best fit. Red squares are the intensity measurements from 3.5, 4.5, 6, and 8 x10^6^ number of cells, fit to the standard curve.

**Supplemental Figure 2:**
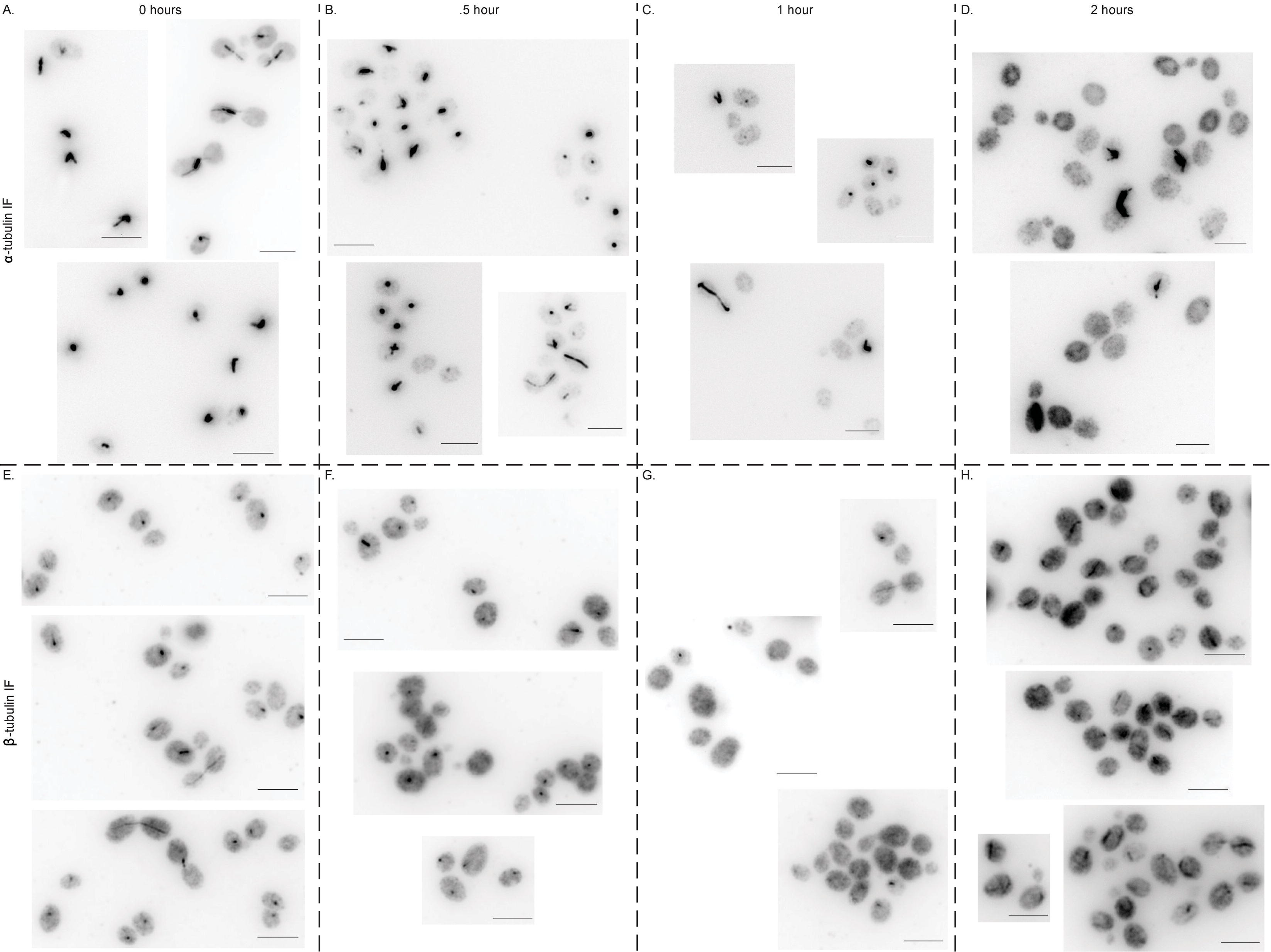
Panel of images through β-tubulin induction. A. Example fields of immunofluorescence images at 0 hr of β-tubulin against α-tubulin. Scale bar = 5 µm. B. Example fields of immunofluorescence images at 0.5 hr of β-tubulin against α-tubulin. Scale bar = 5 µm. C. Example fields of immunofluorescence images at 1 hr of β-tubulin against α-tubulin. Scale bar = 5 µm. D. Example fields of immunofluorescence images at 2 hr of β-tubulin against α-tubulin. Scale bar = 5 µm. E. Example fields of immunofluorescence images at 0 hr of β-tubulin against β-tubulin. Scale bar = 5 µm. F. Example fields of immunofluorescence images at 0.5 hr of β-tubulin against β-tubulin. Scale bar = 5 µm. G. Example fields of immunofluorescence images at 1 hr of β-tubulin against β-tubulin. Scale bar = 5 µm. H. Example fields of immunofluorescence images at 2 hr of β-tubulin against β-tubulin. Scale bar = 5 µm.

**Supplemental Figure 3:**
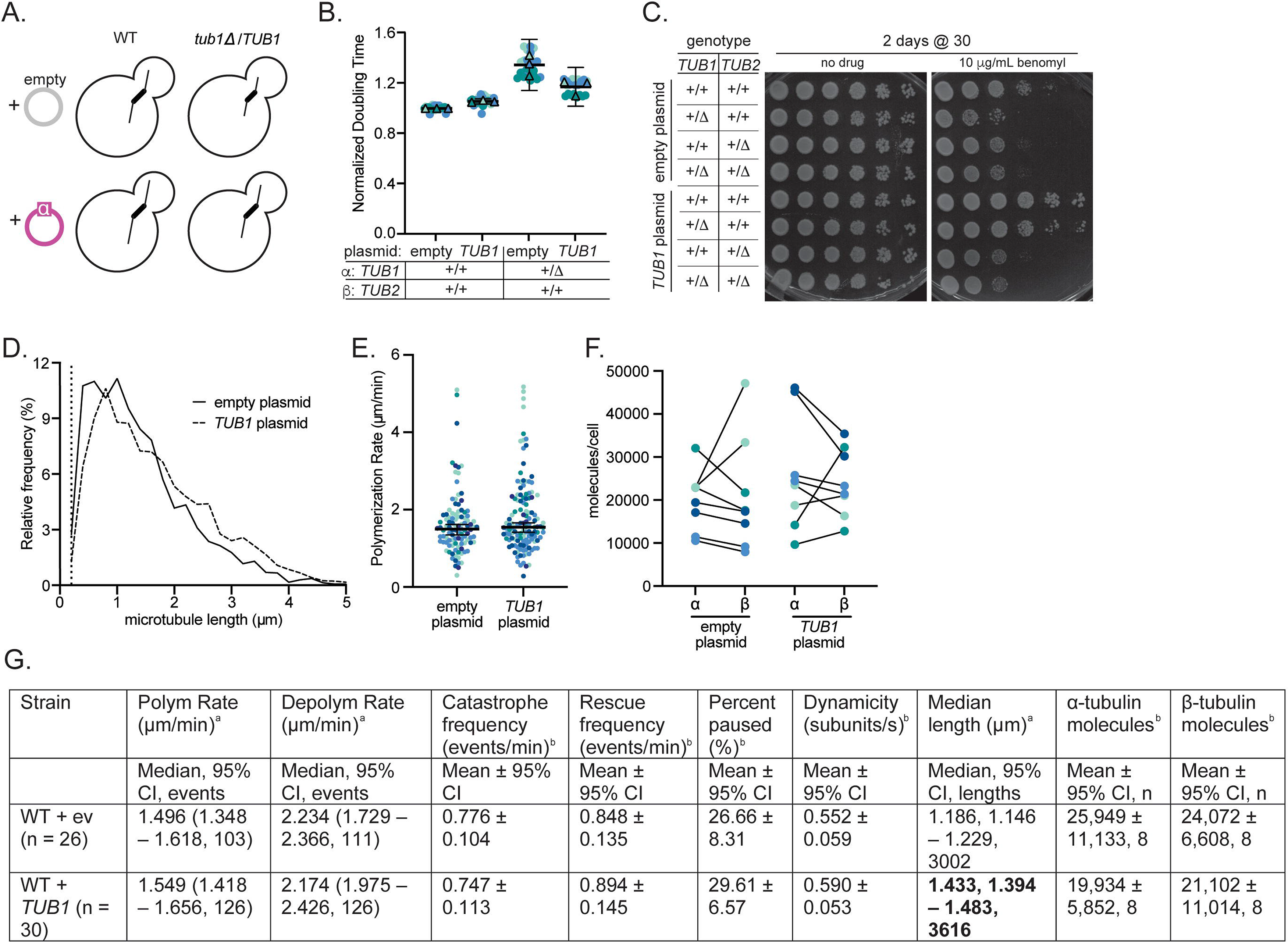
Exogenous copy of *TUB1* does not strongly impact cytoskeleton. A. Schematic of the CEN-based additional copy of *TUB1*. B. Normalized doubling times of wild-type or *tub1Δ/TUB1* with an empty vector or additional copy of *TUB1* on a CEN plasmid. For each genotype, four technical replicates of two biological replicates were used across three independent experiments. Circles represent cultures in separate wells and triangles are the means of each day. Colors indicate independent experiments. Asterisk indicates p< 0.05 between wild type and mutant based on t-test after one-way ANOVA. Bars are mean ± 95% CI. C. Tenfold dilution series of indicated strains were spotted onto rich medium or rich medium supplemented with benomyl. Cells were grown at the indicated temperature for the indicated number of days. D. Histogram of all astral microtubule lengths from time-lapse imaging of wild-type cells with an empty vector or an extra copy of *TUB1*. Data are from at least three separate experiments for each genotype, and a total of at least 26 cells were analyzed for each genotype. E. Polymerization rates of astral microtubules. Each dot represents a single polymerization event and dots are colored by experimental day. Bars are median ± 95% CI. F. Paired molecules of α- or β-tubulin per cell for each genotype. Protein mass (ng) was converted to molecules per cell. Data represent four independent experiments with two biological replicates and each dot is mean molecules per cell of the four technical replicates. Dots are colored by experiment. G. Table of astral microtubule dynamics and soluble tubulin levels in diploid cells +*TUB1* or +ev. Values in bold have p<0.05 from wild type + ev, based on a T-test. ^a^ Median values (95% CI) ^b^ Mean values ± 95% CI ^c^ Mean values ± 95% CI

**Supplemental Figure 4:**
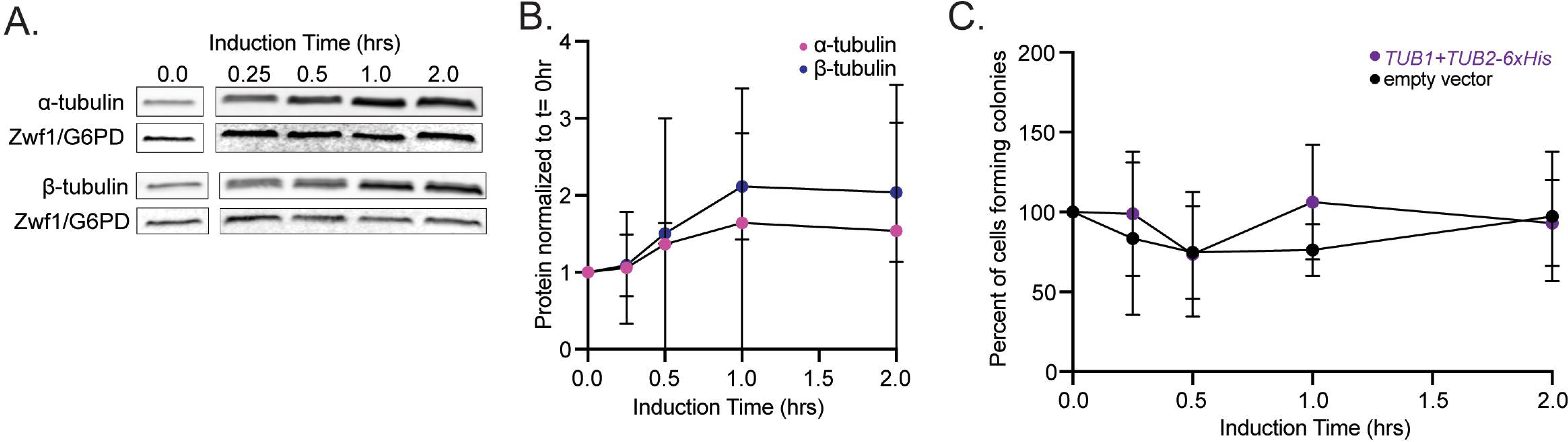
Co-overexpression of α- and β-tubulin is tolerated. A. Representative western blot of β- and α-tubulin during *TUB1* and *TUB2* co-induction. Blots were probed for α- or β-tubulin and Zwf1 (G6PD) as a loading control. B. Quantification of *TUB1* and *TUB2* co-induction for both α- and β-tubulin across overexpression. Dots represent mean ± SD from three independent induction experiments. C. Quantification of cells forming colonies after *TUB1* and *TUB2* induction or with empty vector. Dots represent mean ± SD from three independent induction experiments.

**Supplemental Figure 5:**
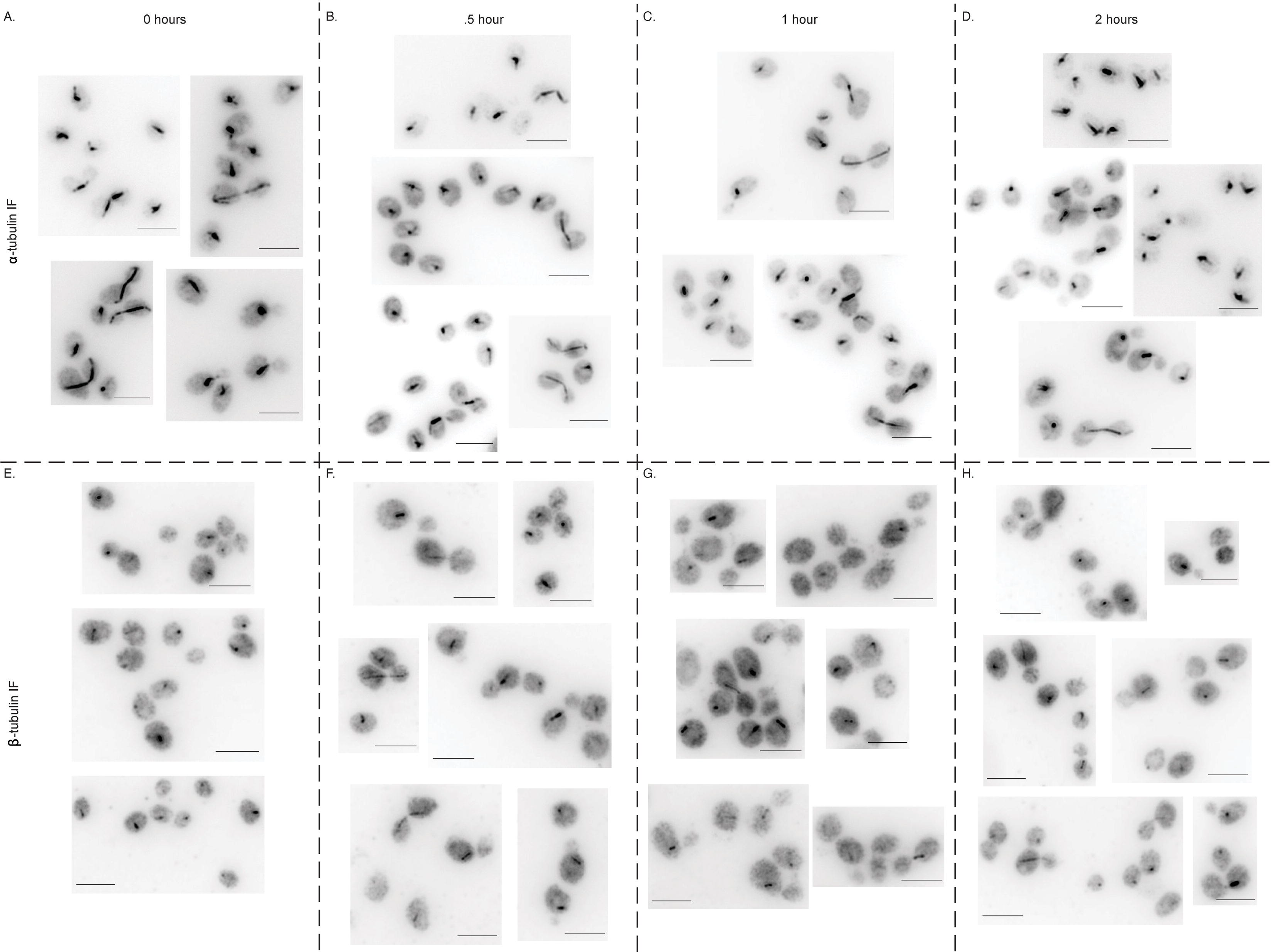
Panel of images through α-tubulin induction. A. Example fields of immunofluorescence images at 0 hr of α-tubulin against α-tubulin. Scale bar = 5 µm. B. Example fields of immunofluorescence images at 0.5 hr of α-tubulin against α-tubulin. Scale bar = 5 µm. C. Example fields of immunofluorescence images at 1 hr of α-tubulin against α-tubulin. Scale bar = 5 µm. D. Example fields of immunofluorescence images at 2 hr of α-tubulin against α-tubulin. Scale bar = 5 µm. E. Example fields of immunofluorescence images at 0 hr of α-tubulin against β-tubulin. Scale bar = 5 µm. F. Example fields of immunofluorescence images at 0.5 hr of α-tubulin against β-tubulin. Scale bar = 5 µm. G. Example fields of immunofluorescence images at 1 hr of α-tubulin against β-tubulin. Scale bar = 5 µm. H. Example fields of immunofluorescence images at 2 hr of α-tubulin against β-tubulin. Scale bar = 5 µm.

**Table S1.**
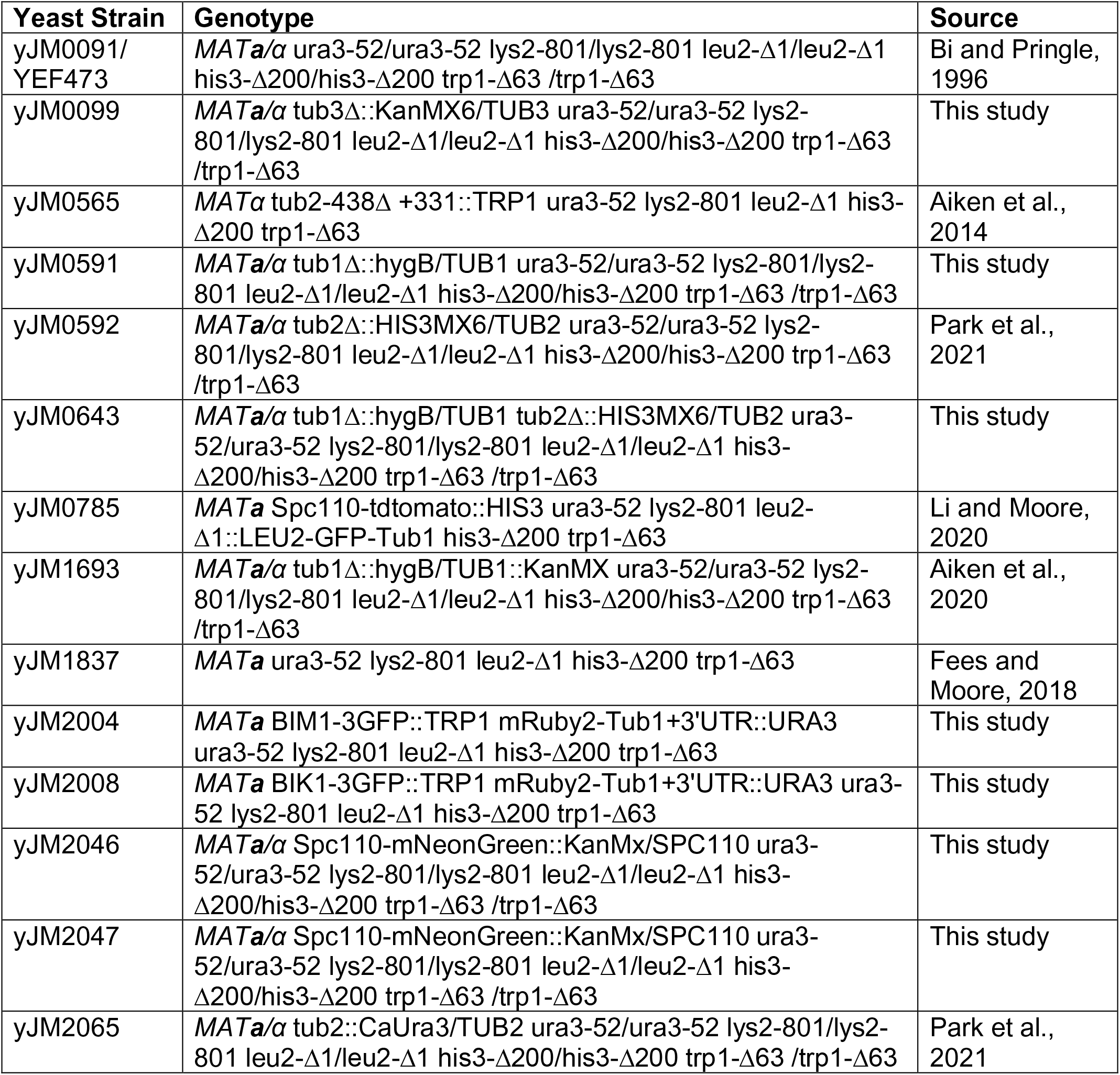
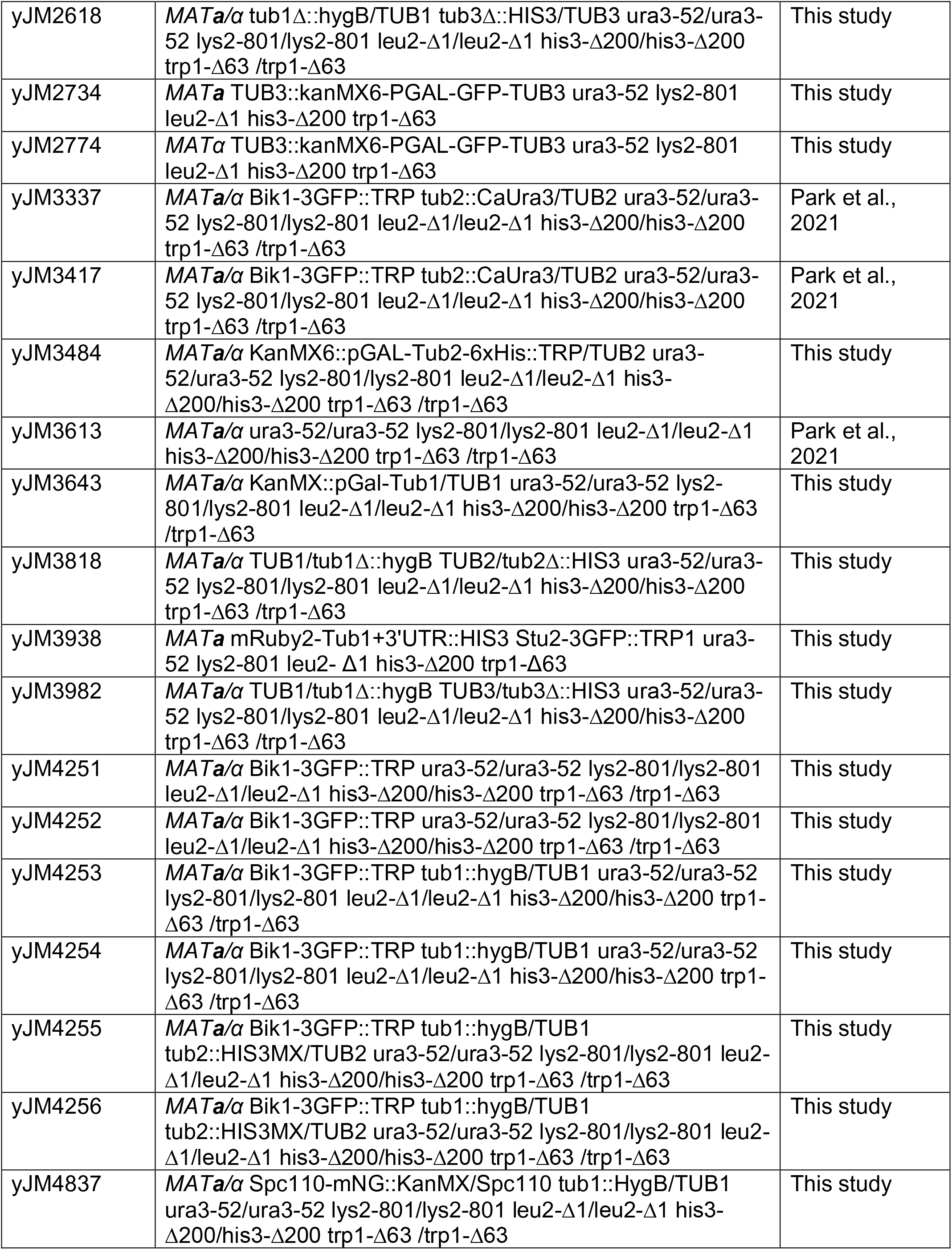
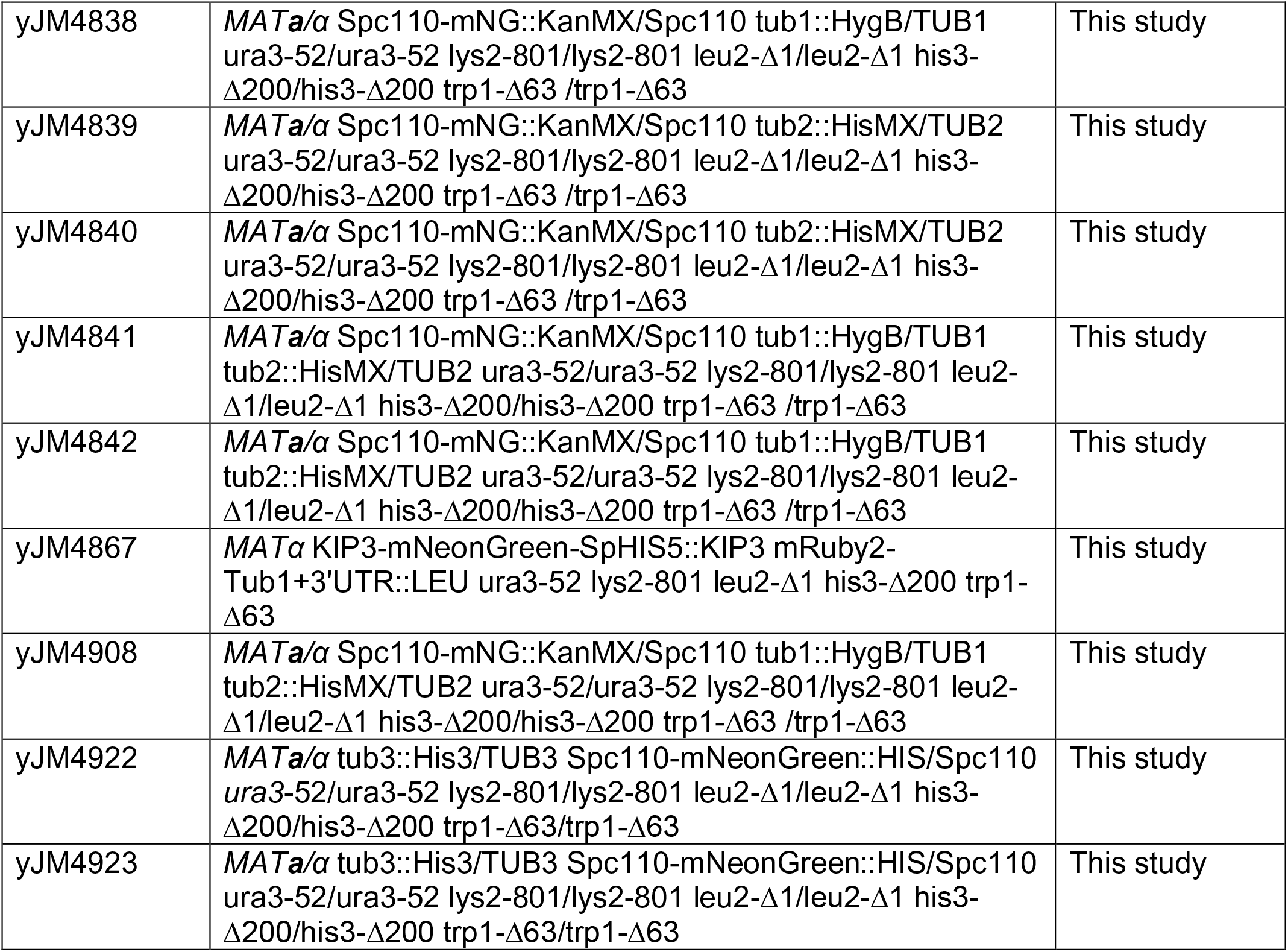
Yeast Strains.

**Table S2.** Plasmids used in this study.

| Plasmid ID | Plasmid Name | Source |
| --- | --- | --- |
| pJM0002 | PB1587 Bik1-3GFP-Trp1 integration plasmid | Carvahlo et al. 2004 |
| pJM0073 | pRS315 | Sikorski and Hieter, 1989 |
| pJM0074 | pRS316 | Sikorski and Hieter, 1989 |
| pJM307 | pURA3:HIS3p:mRuby2-Tub1+3'UTR | Marcus et al., 2015 |
| pJM309 | pLEU2:HIS3p:mRuby2-Tub1+3'UTR | Marcus et al., 2015 |
| pJM310 | pTRP1:HIS3p:mRuby2-Tub1+3'UTR | Marcus et al., 2015 |
| pJM0459 | Stu2-3GFP-TRP integration plasmid | Hoff et al. 2022, preprint |
| pJM0544 | pGAL-Tub2-6xHis::TRP in pRS316 | This study |
| pJM0579 | pGAL-Tub1 in pRS315 | This study |
| pJM0616 | pGAL-Tub2-6xHis::TRP in pRS313 | This study |
| pJM0759 | CEN-Tub1 in pRS316 | This study |

**Table S3.**
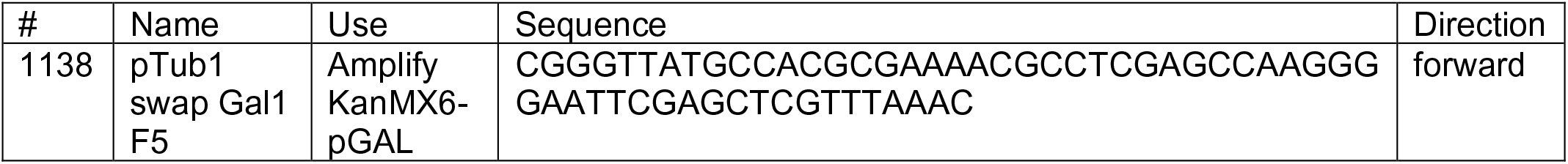
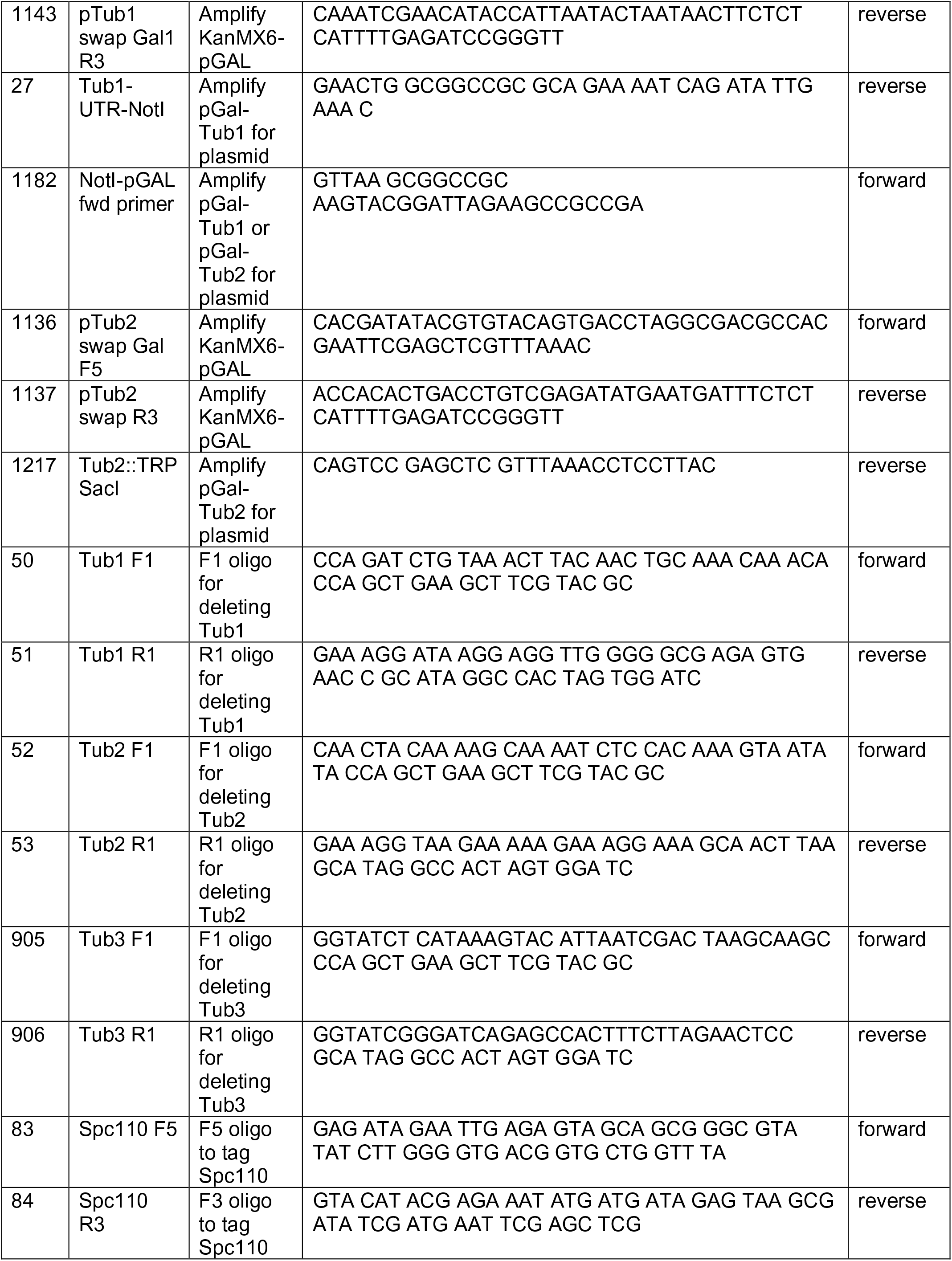
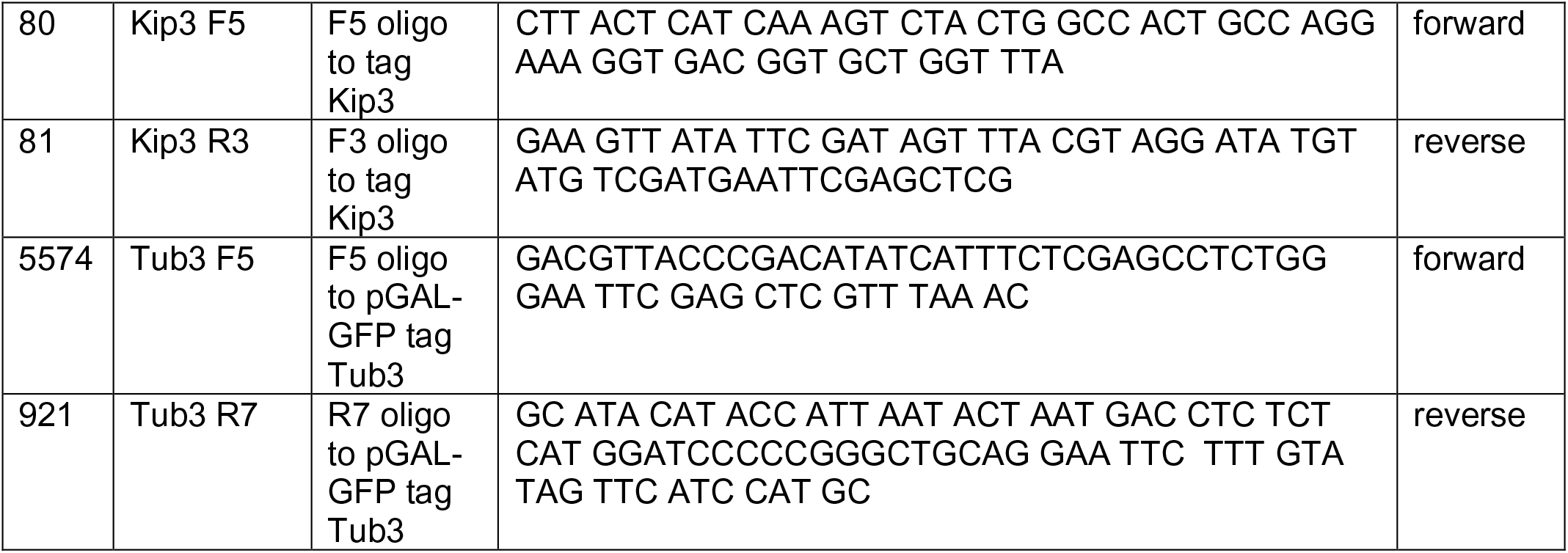
Oligonucleotides used in this study.

## Notes

### Competing Interest Statement

The authors have declared no competing interest.

